# Identification of two distinct phylogenomic lineages and model strains for the understudied cystic fibrosis lung pathogen *Burkholderia multivorans*

**DOI:** 10.1101/2023.01.03.522605

**Authors:** Kasia M. Parfitt, Angharad E. Green, Thomas R. Connor, Daniel R. Neill, Eshwar Mahenthiralingam

**Affiliations:** Cardiff University, Microbiomes, Microbes and Informatics Group, Organisms and Environment Division, School of Biosciences, Cardiff University, CF10 3AX, UK; Department of Clinical Infection, Microbiology and Immunology, Institute of Infection, Veterinary and Ecological Sciences, University of Liverpool, Liverpool, L69 7BE, UK

**Keywords:** *Burkholderia multivorans*, phylogenomics, phenotype, infection modelling, cystic fibrosis

## Abstract

*Burkholderia multivorans* is the dominant *Burkholderia* pathogen recovered from lung infection in people with cystic fibrosis. However, as an understudied pathogen there are knowledge gaps in relation to its population biology, phenotypic traits and useful model strains. A phylogenomic study of *B. multivorans* was undertaken using a total of 283 genomes, of which 73 were sequenced and 49 phenotypically characterized as part of this study. Average nucleotide identity analysis (ANI) and phylogenetic alignment of core genes demonstrated that the *B. multivorans* population separated into two distinct evolutionary clades, defined as lineage 1 (*n* = 58 genomes) and lineage 2 (*n* = 221 genomes). To examine the population biology of *B. multivorans*, a representative subgroup of 77 *B. multivorans* genomes (28 from the reference databases and the 49-novel short-read genome sequences) were selected based on multilocus sequence typing (MLST), isolation source and phylogenetic placement criteria. Comparative genomics was used to identify *B. multivorans* lineage-specific genes: *ghrB_1* in lineage 1, and *glnM_2* in lineage 2, and diagnostic PCRs targeting them successfully developed. Phenotypic analysis of 49 representative *B. multivorans* strains showed considerable variance with the majority of isolates tested being motile and capable of biofilm formation. A striking absence of *B. multivorans* protease activity *in vitro* was observed, but no lineage-specific phenotypic differences demonstrated. Using phylogenomic and phenotypic criteria, three model *B. multivorans* CF strains were identified, BCC0084 (lineage 1), BCC1272 (lineage 2a) and BCC0033 lineage 2b, and their complete genome sequences determined. *B. multivorans* CF strains BCC0033 and BCC0084, and the environmental reference strain, ATCC 17616, were all capable of short-term survival within a murine lung infection model. By mapping the population biology, identifying lineage-specific PCRs and model strains, we provide much needed baseline resources for future studies of *B. multivorans*.

## Introduction

Cystic fibrosis (CF) is a hereditary genetic disorder affecting over 10,500 people in the UK [1]. Mutations in the CF transmembrane conductance regulator gene of people with CF results in several pathological features, with abnormal lung clearance, chronic respiratory infection and severe lung disease being major contributors to morbidity. Although *Pseudomonas aeruginosa* is the most prevalent CF pathogen, *Burkholderia cepacia* complex (Bcc) bacteria, a taxonomic group of closely related *Burkholderia*, emerged as virulent and transmissible CF lung infections in the 1990s [2]. For people with CF, infection with Bcc pathogens can contribute to severe lung function decline and the development of ‘cepacia syndrome’ [3], and those infected also have a lower survival rate after lung transplantation [4]. Whilst the Bcc have been reported at relatively low in prevalence in CF populations (<5%) [4-6], they are of significant clinical consequence because they are hard to eradicate due to their intrinsic resistance to antibiotics, with certain strains being resistant to the 10 most administered antibiotics [7].

*Burkholderia multivorans* is a member of Bcc and is the most isolated *Burkholderia* species in the UK, with 56% of all *Burkholderia* CF lung infection cases (n=361) attributed to the pathogen in 2017 [5]. Earlier surveys of the US showed *B. multivorans* accounted for 37% of *Burkholderia* CF infections at the time [6] and the same dominance was observed in a Canadian study with 45% of 122 *Burkholderia* CF lung infection cases cause by this Bcc species [4]. The epidemiology of *Burkholderia* CF infections also shows that *B. multivorans* has become dominant due to reduced rates of *B. cenocepacia* infection, which is now the second most common Bcc species in multiple CF populations [4-6]. With strict infection control and the resulting absence of patient-patient transmission, the continuing emergence of *B. multivorans* in people with CF suggests that current infections arise sporadically from natural sources such as soil, the rhizosphere and water [2, 6, 8]. However, specific environmental reservoirs of *B. multivorans* remain elusive, with isolates rarely recovered from the natural environment [6, 8].

In contrast to this current epidemiological prevalence of *B. multivorans, B. cenocepacia* has been the most widely studied CF *Burkholderia* [9]. *B. cenocepacia*, is generally considered to be the hyper virulent species within the Bcc [2] and can be separated into two genetic lineages (III-A and III-B) based on the *recA* gene [10, 11]. Recent genomic analysis of *B. cenocepacia* provided further evidence to show that the species should be split into at least two different species based on average nucleotide identity (ANI) differences [12]. The latter studied argued for the name *“Burkholderia servocepacia”* to be attributed to strains falling into the *recA* III-B grouping, but this proposition was invalid based on taxonomic and naming criteria. *Burkholderia orbicola* sp. nov. [13] has now been validly proposed as the species name for the genomic taxa represented by *“Burkholderia servocepacia*.” Overall, multiple studies have shown that epidemic and transmissible CF strains can be found in both *B. cenocepacia* and *B. orbicola* sp. nov. [9]. For example, *B. cenocepacia* III-A strains are associated with poor clinical outcome and major morbidity in several CF populations [2, 6], with the ET-12 strain being notable in virulence and prevalence, together with multiple other intercontinentally dispersed multilocus sequence types (MLST) [9]. Virulence factors such as the cable pilus, cenocepacia pathogenicity island and multiple quorum sensing-dependent pathogenicity traits, have also been characterised for *B. cenocepacia* [9].

In comparison, much less is known about the pathogenicity of *B. multivorans* in CF. The presence of non-mucoid isolates of Bcc bacteria have been shown to be correlated with greater decrease in lung function of infected individuals [14], and this mucoid variation in *B. multivorans* was associated with changes in metabolism, motility, biofilm formation and virulence [15]. Within-strain genomic evolution has been studied for multiple isolates recovered over 20 years from an individual with CF [16]. The average evolutionary substitution mutation rate for this single *B. multivorans* strain was low overall, at 2.4 mutations per year, with one intra-strain lineage evolving more rapidly than the others through non-synonymous mutations [16]. Alterations in the *B. multivorans* phenotype during chronic infection were linked to mutational changes in antimicrobial resistance, biofilm formation and LPS O-antigen presentation gene pathways [16]. Another study obtained genome sequences from 111 clonal isolates of *B. multivorans* from a single person with CF, as their lung disease progressed [17]. Statistically significant accumulations of mutations in loci contributing to increased antimicrobial resistance were seen in this single-strain evolutionary study [17]. Genomic comparison of *B. multivorans* isolates isogenic by MLST, but from CF infection and natural environmental sources, demonstrated that the same genomic lineages occur in these different niches and across different continents [18].

A comparison of multiple genetically distinct *B. multivorans* strains that includes both phenotypic and genomic characterisation of the species has not yet been made. Our study aimed to unpick the phylogenomics and basic pathobiology of *B. multivorans*, as both a species and an understudied CF lung pathogen. Whole Genome Sequencing (WGS) was used to characterise 73 genetically diverse *B. multivorans* strains drawn from multiple sources, MLST strain types, and geographic regions. A further 210 *B. multivorans* genomes were obtained from publicly accessible databases and analysed phylogenetically. Twenty-eight of the database sequences were combined with 49 of the *de novo* genome sequenced strains to produce a representative strain panel (*n* = 77). The *B. multivorans* strain panel encompassed 61 unique MLST sequence types (STs; 5 novel), focussing on CF isolates (*n =* 60) and including strains from the environment (*n* = 8), non-CF infection (*n* = 8), and one isolate of an undetermined source. The phenotypic features of 49 representative strains selected from this panel were investigated by swimming and swarming motilities, biofilm formation, exopolysaccharide production and protease production, and three strains were also tested for survival in a mammalian respiratory inhalation lung infection model [19, 20]. From this analysis, an evolutionary split into 2 genetic lineages was shown for *B. multivorans* as a CF pathogen.

## Methods

### Bacterial strains and incubation conditions

The bacterial strains phenotypically studied, and genome sequenced in this study were drawn from the *Burkholderia* strain collection at Cardiff University and additional recognised strain repositories [20, 21] (Table 1). A complete list of the 283 isolates and their genomes analysed within the study is provided in Supplementary Table S1. The isolates studied were recovered from a range of sources including CF, Chronic Granulomatous Disease (CGD), non-CF clinical infections (NON-CF), the natural environment (ENV) and healthcare environments (ENVH). Stock cultures were stored at -80°C in cryogenic vials by resuspension of fresh growth in Tryptic Soya Broth (TSB; Oxoid) containing 8% (v/v) dimethyl sulfoxide (Sigma Aldrich). Culture purity was determined by plating frozen stocks onto Tryptic Soy Agar (TSA) (Oxoid) and incubating plates for 24 - 48 h at 37°C. Overnight cultures were made by taking a swab from a fresh TSA plate and transferring into 3 ml of TSB. Cultures were grown for 18-20 h at 37°C using continuous shaking on a rotating platform set to 150 r.p.m.

**Table 1.**
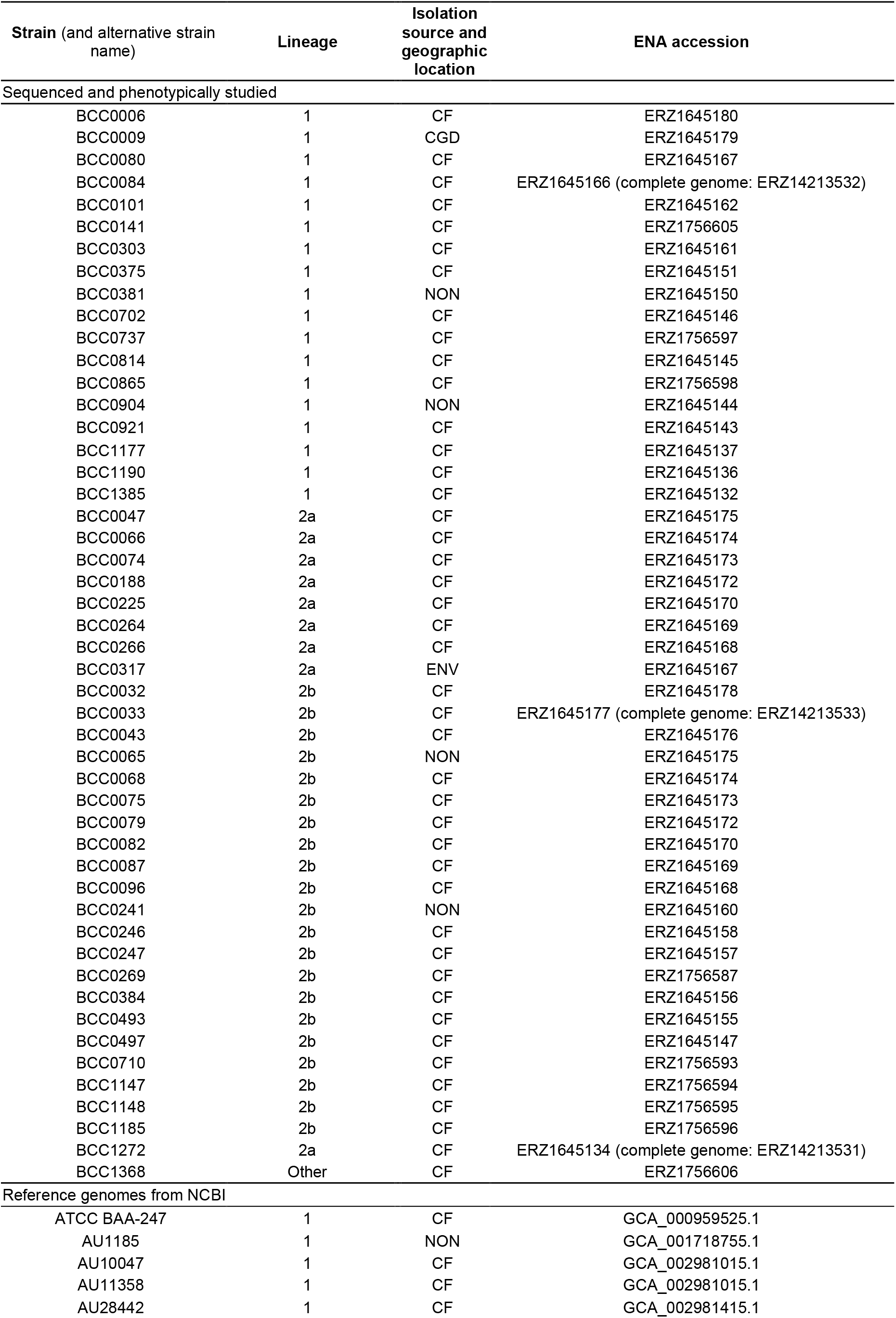
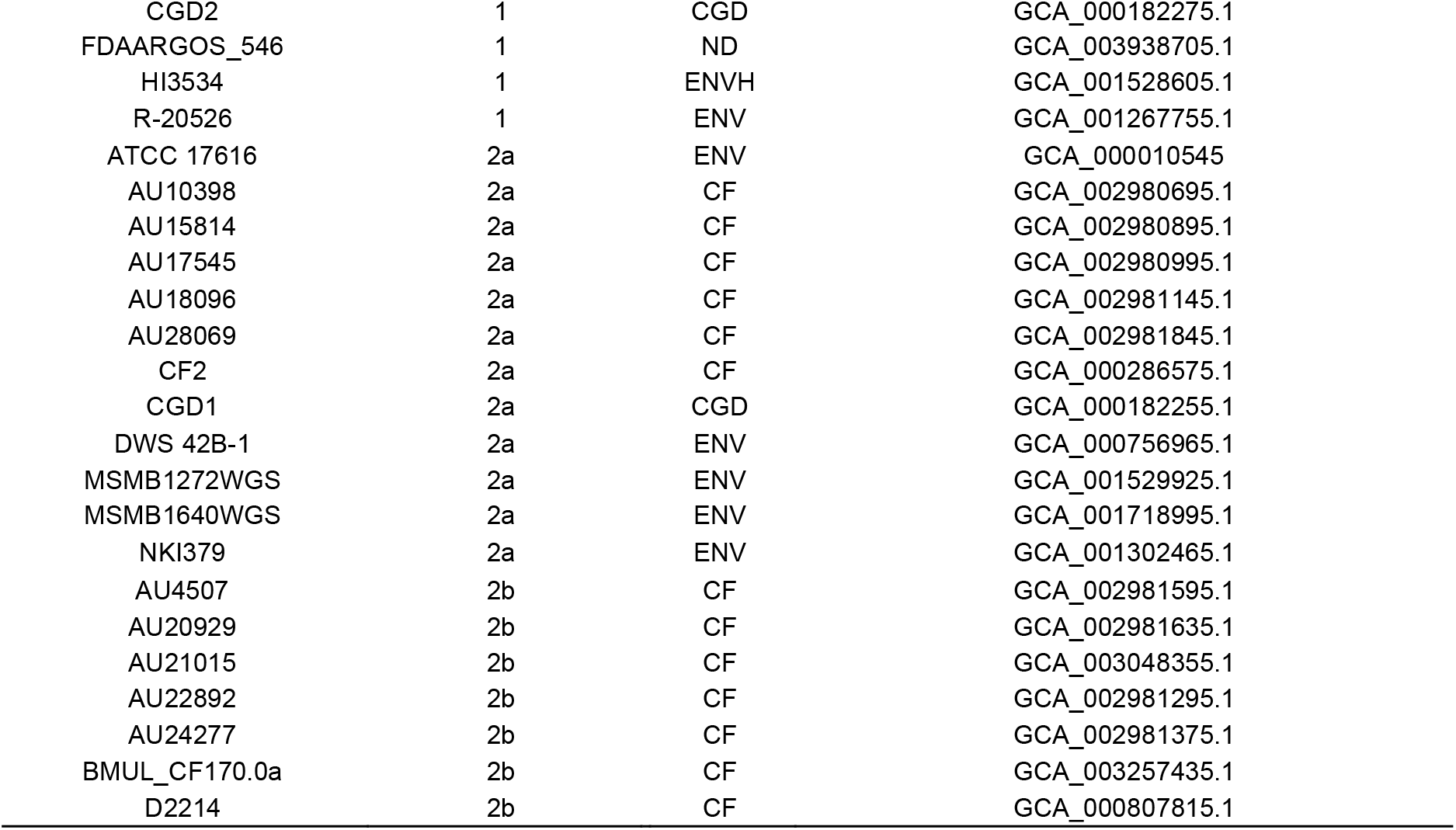
The selected *B. multivorans* strain panel (*n = 77*) including 49 phenotypically characterized strains sequenced in this study.

### Genome sequencing of *B. multivorans*

*B. multivorans* strains for genome sequence were selected based on their source, geographic distribution, and MLST-based genetic diversity [22, 23] (Table 1 and Table S1). After revival and purity checking, 3 ml overnight cultures were subjected to DNA extraction using an automated Maxwell® 16 Tissue DNA purification kit and following the manufacturer’s instructions (Promega, UK). For long-read complete genome analysis, DNA was extracted using a DNA Wizard Kit (Promega, UK). Upon extraction, each DNA sample was transferred into non-stick 1.5 ml microtubes and stored at -20°C. DNA samples were checked for purity using the *B. multivorans* specific *recA* primers, BCRBM1 and BCRBM2 [10], with PCR amplicons visualised on a 1.5% (w/v) agarose gels, prior to Sanger sequence analysis to confirm they were *B. multivorans*.

A total of 73 *B. multivorans* strains were subjected to short-read WGS using an Illumina MiSeq V2 platform within the Genome Hub at Cardiff School of Biosciences. Genomic reads were assembled and annotated using the shared Cloud Infrastructure for Microbial Genomics (CLIMB) computing facility [24]. Illumina reads were subjected to the Trim Galore v0.4.4 [25] wrapper script. This utilises Cutadapt v1.9.1 [26] for automated quality and adapter trimming and FastQC v0.11.4 [27] for quality control. MultiQC v1.7 [28] Python package was used to compile a single file report and interactive report for the samples, helping to streamline quality control screening. All genomes possessed sufficient quality to take forward for phylogenomic analyses (Table S2).

To assemble the bacterial genomes, we used the Unicycler v0.4.7 [29] assembly pipeline, which utilizes SPAdes [30] for optimizing and streamlining *de novo* assembly of the genome contigs. Complete genome sequence analysis was performed for the three selected model strains (BCC0033, BCC0084 and BCC1272) using long read PacBio technology (carried out by Novogene, UK). The PacBio FASTQ reads were subjected to the Trycycler pipeline (v0.4.1) [29] and provide complete assemblies of four contigs (the 3 genomic replicons and a large plasmid in each strain). DNA sequence reads from the selected database genomes were also re-assembled and all 283 *B. multivorans* genomes were subjected to Prokka v1.14.0 [31] to annotate the sequences and provide provides output files suitable for phylogenomic analysis. Accession numbers for the genome sequences obtained in this study are provided in Table 1.

### Genomic taxonomy, phylogenomic and MLST analyis

To confirm the taxonomic identity of the *B. multivorans* genomes and filter out contaminating DNA, the Minikraken database from Kraken2 v2.08-beta [32] was used. QUAST v5.0 [33] was used to assess quality and respective statistics for the genomic assemblies. To confirm species taxonomy, the pairwise ANI was calculated for the *B. multivorans* genomes using the Python3 module and script PyANI v0.2.9 [34]. A 95% threshold was used as an accepted standard to confirm that all strains were the same species in accordance with the Genomic Taxonomy database [35] and recent taxonomic analysis of *Burkholderia* genomes [36].

Phylogenomic and pan genome analysis was performed as follows. The GFF annotated genome-file outputs from Prokka [31] were evaluated in the Roary v3.12.0 pan genome pipeline [37] to assess the core and accessory genome of all 283 *B. multivorans* genomes. The command was performed using the default settings. MAFFT [38] was used to create the Roary core gene alignment output file. Phylogenetic trees were built using maximum likelihood (GTRGAMMA model) Randomized Accelerated Maximum Likelihood (RAxML v8) [39], supported by 100 bootstraps. The *B. dolosa* AU0158 complete genome was initially used to root phylogenetic trees as a closely related Bcc species; subsequent trees were rooted with the *B. multivorans* BCC1638 genome (Table 1). Sequence types were determined for all *B. multivorans* strains using MLSTcheck, utilizing PubMLST blast schemes [40] (Table 2).

**Table 2.**
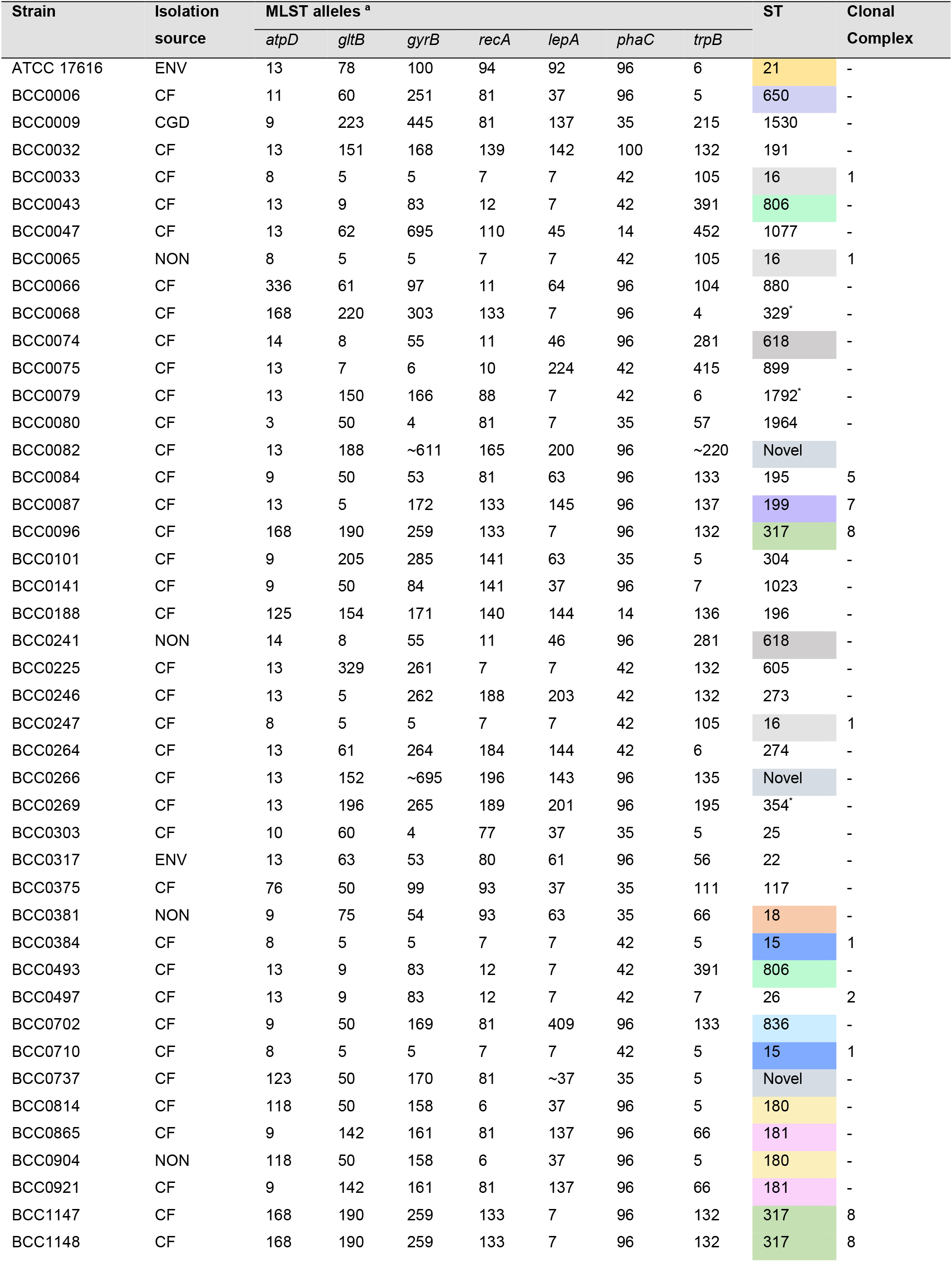
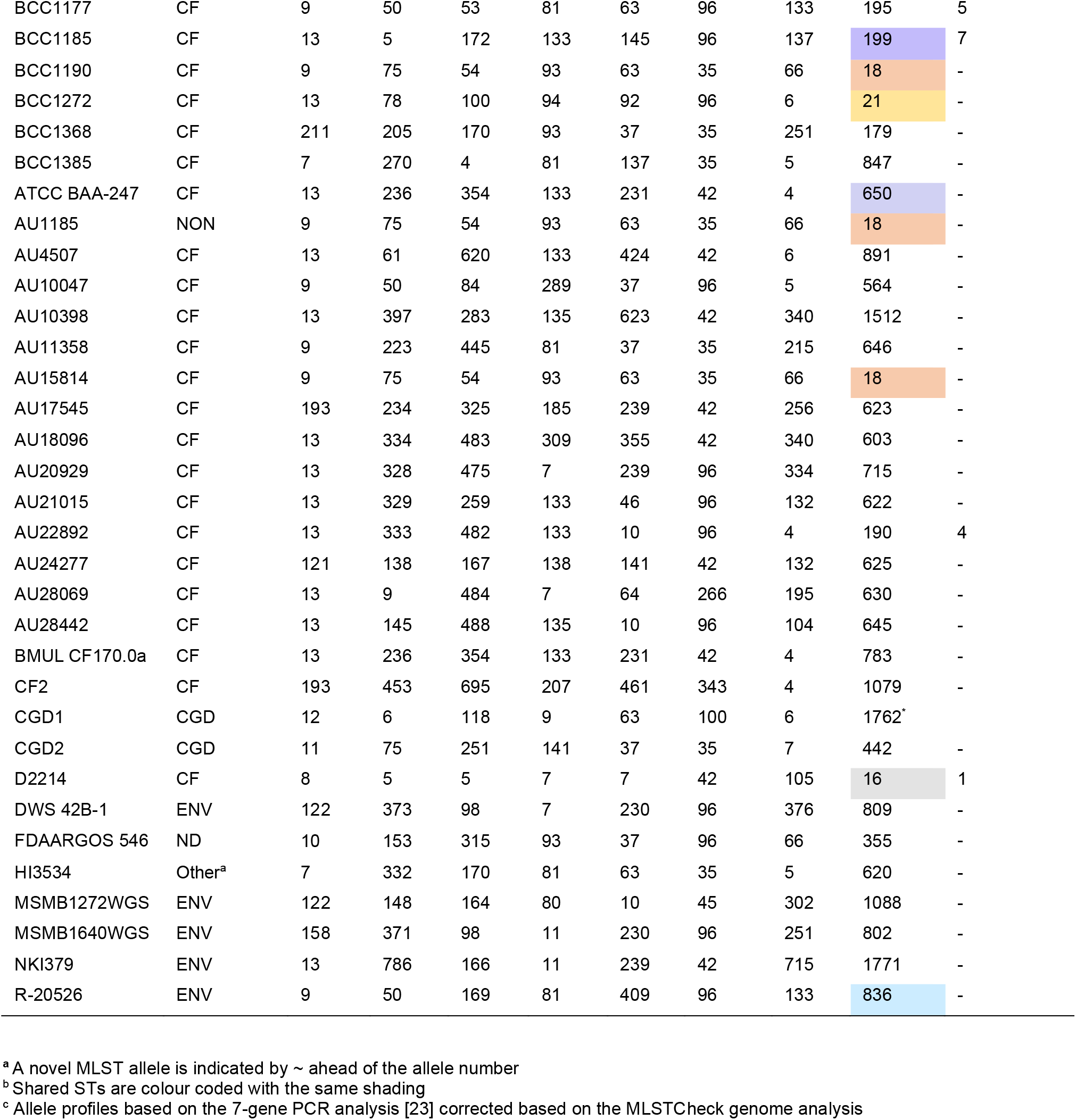
MLST alleles and Sequence Type (ST) for the 77 strain *B. multivorans* panel.

### Assessment of swimming and swarming motilities

Motility of *B. multivorans* was measured using a modified method from Rashid and Kornberg [41]. Agar plates were prepared and dried on an even surface 24 hours before use to ensure consistent moisture content, with each plate containing 20 ml. Agar concentrations were made using 0.3% (w/v) LB for swimming assays and 0.5% (w/v) LB and 0.5% (w/v) basal salts medium supplemented with 0.4% (w/v) glucose (BSM-G) for swarming assays. Swimming motility was assessed by inoculating the agar, through to the base, with a sterile toothpick. Swarming motility was assessed by surface inoculation with a sterile toothpick. Plates were inverted and wrapped in sealed Petri-dish bags to prevent drying. Plates were incubated for 37°C and zones were measured at 24 h, averaging two perpendicular measurements. Each isolate was assigned a category: non-motile ≤ 5 mm, low motility 5-25 mm, intermediate motility 25-50 mm, and high motility ≥ 50.0 mm.

### Biofilm formation of *B. multivorans*

A crystal violet and 96-well PVC plate growth assay [42] was used to determine the biofilm mass formation of *B. multivorans* isolates. Overnight cultures were diluted to roughly 10^5^ c.f.u ml^-1^ in TSB in Falcon tubes. These were gently mixed using a vortex before transferring 100 µl into 96-well plates. The outer wells were left empty to prevent drying and *B. multivorans* biofilms left to form over 24 h by static incubation of the plates at 37°C. After removal of growth media and washing as described [42], biofilm biomass was stained with a solution of 0.1% (w/v) crystal violet for 20 mins. The plates were washed, allowed to dry and the absorbance at 570 nm read for a 200 µl solubilization of the biomass stain in each well using 70% ethanol.

### Growth rate of *B. multivorans*

A Bioscreen C instrument (Labsystems, Finland) was used to determine the bacterial growth dynamics of *B. multivorans* isolates. Cultures (200 µl in TSB) were inoculated with approximately 10^6^ c.f.u. ml^-1^ using an optical density-based standardization of fresh overnight liquid growth. Growth was monitored over 48 h with incubation at 37°C. Well absorbance readings using a wideband filter (450-580 nm) were performed every 15 minutes after 10 seconds of medium shaking. A scatterplot analysis was performed in Microsoft Excel to visualize the growth curves. The data was further analysed using the GcFit function of the grofit package [43] which utilizes R statistical software [44] to output specific parameters of lag phase, maximum growth rate and maximum culture density.

### Exopolysaccharide and protease production by *B. multivorans*

Exopolysaccharide (EPS) production of the *B. multivorans* strains was determined using yeast extract medium (YEM) agar as described by Zlosnik, Hird [45]. The original protocol was used for the agar preparation, with no adaptations. *B. multivorans* was streaked for single colonies from freezer stocks onto the agar plates before incubating for 48 hours at 37 °C. EPS was visually categorized into the following five groups based on the literature [45]: - (non-EPS producing), + (partially mucoid), ++ (low mucoidicity), +++ (medium-high mucoidicity) and ++++ (very high mucoidicity). *B. multivorans* protease production was assessed using a modified protocol from Morris, Evans [46]. The lactose-free skimmed-milk agar was prepared as per the original protocol. Overnight cultures were diluted to ∼10^7^ c.f.u. ml^-1^. Aliquots of 10 µl culture were placed onto the protease media in triplicate. Plates were left to completely dry before being inverted and incubated at 37°C for 24 h. Protease production was measured by taking the average of two perpendicular measurements of resulting colony and the zone of clearing around it (mm). A final protease production value was obtained by subtraction of colony size from the zone of clearing. *P. aeruginosa* LESB58 was used as a positive control for every protease assay.

### Construction of *B. multivorans* fluorescent reporter strains

Electroporation was used to introduce the plasmid vector pIN301-eGFP [47] into the selected *B. multivorans* model strains as follows. Overnight cultures of *B. multivorans* (strains BCC0033, BCC0084, BCC1272, and ATCC 17616) were grown in TSB. These were diluted to an OD_600 nm_ 0.1 (∼10^7^ c.f.u. ml^-1^) in 3 ml TSB before incubating for approximately 4 hours at 37°C, shaking at 150 rpm. This incubation step enabled the *B. multivorans* cultures to reach OD_600nm_ of ∼1 and a 2 ml aliquot of culture was spun down in a centrifuge for 5 minutes at 4000 r.p.m. The pellet was twice washed with 2 ml sterile ddH_2_O before re-suspending 30 µl of ddH_2_O. 10 ng of room temperature pIN301-eGFP DNA was added to the suspension, and the suspension transferred to sterile 2 mm electroporation cuvette (Thermo Fisher). After electroporation using 2500V, with a field capacity of 12.5 kV/cm, 1 ml sterile TSB was used to recover the electroporated cells for 1 hour at 37°C with shaking at 150 r.p.m. The revived cultures were plated on TSA supplemented with 50 µg/ml chloramphenicol and incubated for 24 h at 37°C before examining under UV light to confirm eGFP::pIN301 plasmid uptake. To confirm that the eGFP::pIN301 derivative was the same as the parental strain, genotyping using by Random Amplification of Polymorphic DNA (RAPD) PCR and primers 270 and 272, was performed as described [48].

### *B. multivorans* lineage-specific PCR primer design

A pan-genome wide association study (GWAS) approach [49] against gene presence-absence output file determined via Roary analysis [37] was used to identify genes unique to each lineage. Four target genes were identified, and PCR primers designed for each as follows (Table S3). The genes were extracted from the *B. multivorans* strain panel genomes (Table 1) using Bedtools [50] and aligned using MAFFT [38]. Regions of within-lineage similarity were selected for primer design, and the resulting primer sequences checked for basic specificity using NCBI primer BLAST, and hairpin structures using the Oligoanalyzer tool (Integrated DNA Technologies). Forward and reverse primers for each gene, together with their genomic location are provided in Table S3; the information of the PCRs selected for testing are provided in Table 3. The PCR primers were synthesised (Eurofins Genomics) and optimised on the 4 *B. multivorans* model strains using a gradient PCR. Thermal cycling conditions of an initial denaturation (95°C, 5 min), 30 cycles of denaturation (95°C, 30 s), annealing (30 s; see Table 3 for temperature) and extension (72°C, 30 s), followed by a final extension (72°C, 10 min) were used. The PCRs were evaluated on the DNA from the 49 phenotypically characterised *B. multivorans* strain panel isolates (Table 1), with *B. ambifaria* and *B. cenocepacia* DNA used as negative controls. PCR products were separated by electrophoresis on a 1.2% agarose gel and visualised using a UV transilluminator.

**Table 3.**
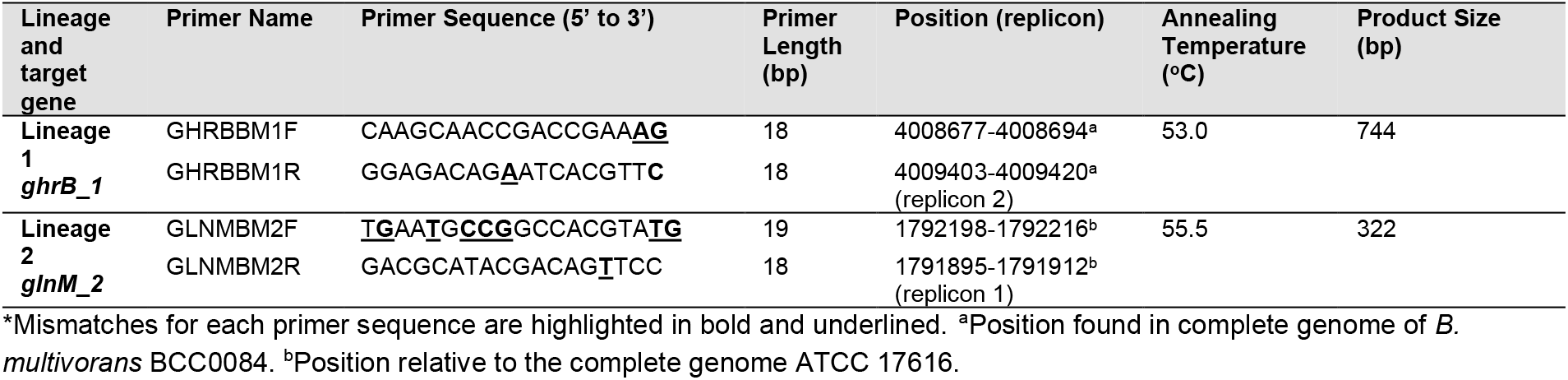
Lineage-specific *B. multivorans* target genes and PCR primer sequences.

### Murine lung infection modelling

A murine chronic lung infection model successfully applied to *P. aeruginosa* [19, 51] and *Burkholderia ambifaria* [20] was used to evaluate basic infection traits of 3 model *B. multivorans* strains. These included wild-type BCC0033 and ATCC 17616, and GFP-tagged derivatives BCC0084 eGFP::pIN301 and BCC0033 eGFP::pIN301. Murine infection modelling was performed with full ethical permission at University of Liverpool under project licence PP2072053 (approved by the UK home office and the University of Liverpool Animal Welfare and Ethical Review Board). BALB/c female 6–8-week-old mice (Charles River, Margate, UK), were used for all experiments and randomly assigned to a cage of four mice by staff independent of the study. Mice were then housed in individually-ventilated cages for 7 days before *B. multivorans* infection, to allow acclimatisation. Overnight cultures of each *B. multivorans* strain were grown in TSB using a single colony inoculation, and subcultured in fresh TSB supplemented with 20% foetal bovine serum (FBS) for ∼6 hours to allow them to reach mid-exponential phase. Standardised suspensions of each *B. multivorans* strain were prepared, plated to determine viability, and were stored at -80°C.

Murine infections were performed using a protocol from Green *et al*. [52], whereby the frozen *B. multivorans* stock suspensions were thawed at room temperature, were harvested by centrifugation and resuspended in phosphate-buffered saline (PBS). For each *B. multivorans* strain, 24 mice were intranasally infected with ∼10^7^ c.f.u ml^-1^ within a 50 µl suspension. This was performed under light anaesthesia using O2/isoflurane. The nasopharynx and lungs were removed, post-mortem at 1-, 3-, and 5-days post-infection, before homogenizing in 2 ml sterile PBS using a hand-held tissue homogeniser (VWR). Ten-fold serial dilutions of tissue homogenates were then prepared and plated onto *B. cepacia* selective agar (BCSA) (Oxoid, UK). *B. multivorans* viable cell counts were enumerated after incubation for up to 48 h at 37°C. For each infection strain, the isolates at day 3 and 5-post infection were pooled from the 8 mouse replicate plates into one stock for the nasopharynx and one for the lungs. Genomic DNA was extracted from the post-infection isolate pools as described above and subjected to short-read Illumina sequencing (Novogene; Cambridge, UK). Genome sequences were then checked for quality and assembled as above. Snippy V3.2-dev was then used for SNP analysis [53].

### Statistical analysis

The phenotypic analysis experiments were performed as 3 biological replicates unless stated otherwise. All statistical analysis was performed in R [44]. The data generated from the analyses within the study was considered to have non-normal distribution, hence, the Kruskal-Wallis chi-squared test (2 comparisons) or Dunn Test with Benjamini-Hochberg correction (3 or more comparisons) were used for statistical evaluation as stated.

## Results

### De novo genomic analysis of *B. multivorans* as a Bcc species

A total of 73 *B. multivorans* genomes were short-read sequenced as part of this study (49 shown in Table 1; additional strains in Table S2) and all possessed high quality draft genome sequences (Table S2). The assembled contigs produced genomes which ranged in size from 6.02 Mb to 7.1 Mb, with an average of 6.514 Mb and mean G+C content of 67.14%. The number of predicted coding sequences (CDS) ranged between 5975 and 7374 CDS, and between 43 and 67 RNA encoding loci were identified per genome (Table S2). When the 73 strain genomes were combined with publicly available sequences to form the 283 master genome panel (Table S1), the genome metrics remained consistent with a mean GC content of 67.04%, sequence length of 6.5 Mb, N50 value of 338304, and mean CDS of 5814 found for *B. multivorans*.

*Burkholderiales* taxonomy has been extensively reclassified and continues to expand in terms of novel taxa. For example, recent phylogenomic analysis of 7 *Burkholderiales* genus clades (*Burkholderia, Paraburkholderia, Trinickia, Caballeronia, Mycetohabitans, Robbsia*, and *Pararobbsia*) predicted that 235 genomic species groups existed within a set of 4000+ genomes that encompassed 129 validly named species [36]. To gain insights into the *B. multivorans* species population biology and confirm the taxonomic classification of strains, ANI analysis was used as the current gold-standard in bacterial genomic taxonomy [35]. Analysis was initially performed on the large dataset of 283 *B. multivorans* genomes (Table S1), with a sub-set of 77 strains representative of the genomic diversity selected for further analysis (Table 1; environmental, *n* = 8; non-CF infection, *n* = 8; CF, *n* = 60; and 1 undetermined source).

Using the species threshold of 95% ANI [54] which has also proven appropriate for the majority of *Burkholderia* sensu lato genomic species [36], the *B. multivorans* isolate genomes (all 283 and the 77 strain panel) comprised a single genomic taxa (Figure 1). The mean ANI for the 77 *B. multivorans* examined was 98.59% and ranged from 97.24% to 100.00% identity. An ANI heatmap of the 77 strains demonstrated the presence of two prominent groups within the *B. multivorans* population that had further evolved towards more restricted identity (Figure 1). These were designated ANI group 1 (*n =* 28; mean ANI of 99%) and ANI group 2 (*n* = 49; mean ANI of 98%). Further ANI sub-groupings were apparent within ANI group 2, designated 2a and 2b. The *B. multivorans* CF strain BCC1368 formed an outlying ANI group and was designated as “other,” but was still above the 95% ANI threshold of the species (Figure 1).

**Figure 1.**
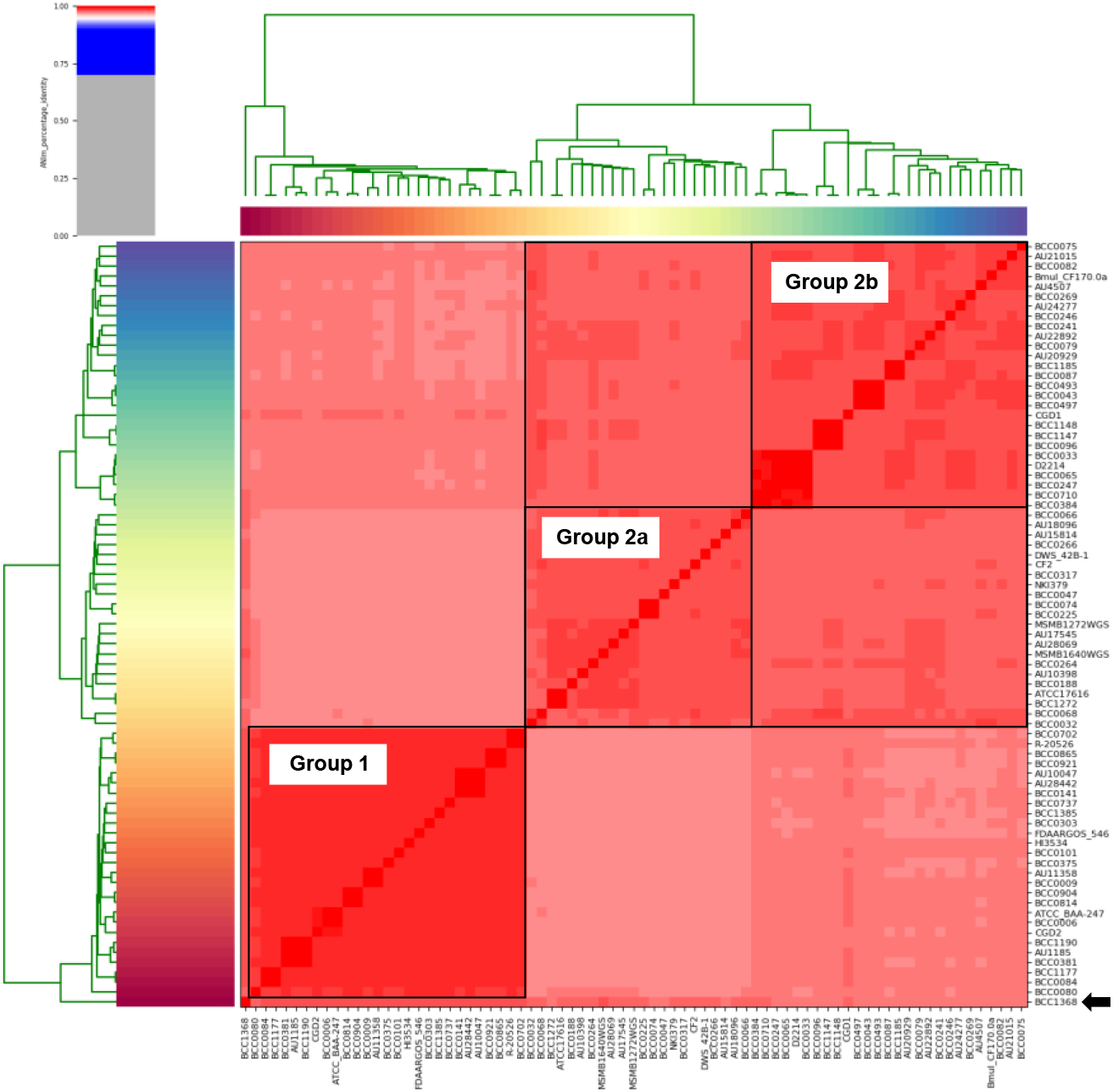
*B. multivorans* is a single genomic species comprised of 2 major ANI sub-groups. An ANI heatmap of the 77 sub-selected *B. multivorans* strains was generated using the PyANI. The ANI percentage identity scale is shown (top left) with all red regions >95% identity. The two major ANI groups, 1 and 2, are indicated with the further 2a and 2b sub-groups labelled. The outlier strain BCC1368 is indicated as by the black arrow (bottom right) and was still >95% ANI in terms of similarity with the other *B. multivorans* genomes (bottom right).

### Core gene phylogenomics corroborates that *B. multivorans* has two major evolutionary lineages

To reconcile an evolutionary basis for the *B. multivorans* ANI population biology (Figure 1), core gene phylogenies were analysed (Figure 2). A master phylogeny was created from the 283 *B. multivorans* genomes using RAxML v8 [39] and alignment of 4319 core genes present in all samples (Figure 2). The phylogenomic tree confirmed the *B. multivorans* population structure was comprised of two major evolutionary lineages, with the greatest diversity and further sub-groupings apparent in lineage 2. The isolate source distribution for the 283 genomes was as follows (Table S1): CF, *n* = 248; CGD, *n* = 6; non-CF clinical infection (*n* = 11); ENV, *n* = 23; ENVH, *n* = 1 and isolates of unknown source, *n* = 2. CF strains were distributed throughout the phylogeny, with lineage 2 containing the majority of the CF strains (n=193) compared to lineage 1 (*n* = 45); four CF strains, including the BCC1368 ANI outlier (Figure 1), clustered within the ‘other’ *B. multivorans* lineage (Figure 2a).

**Figure 2.**
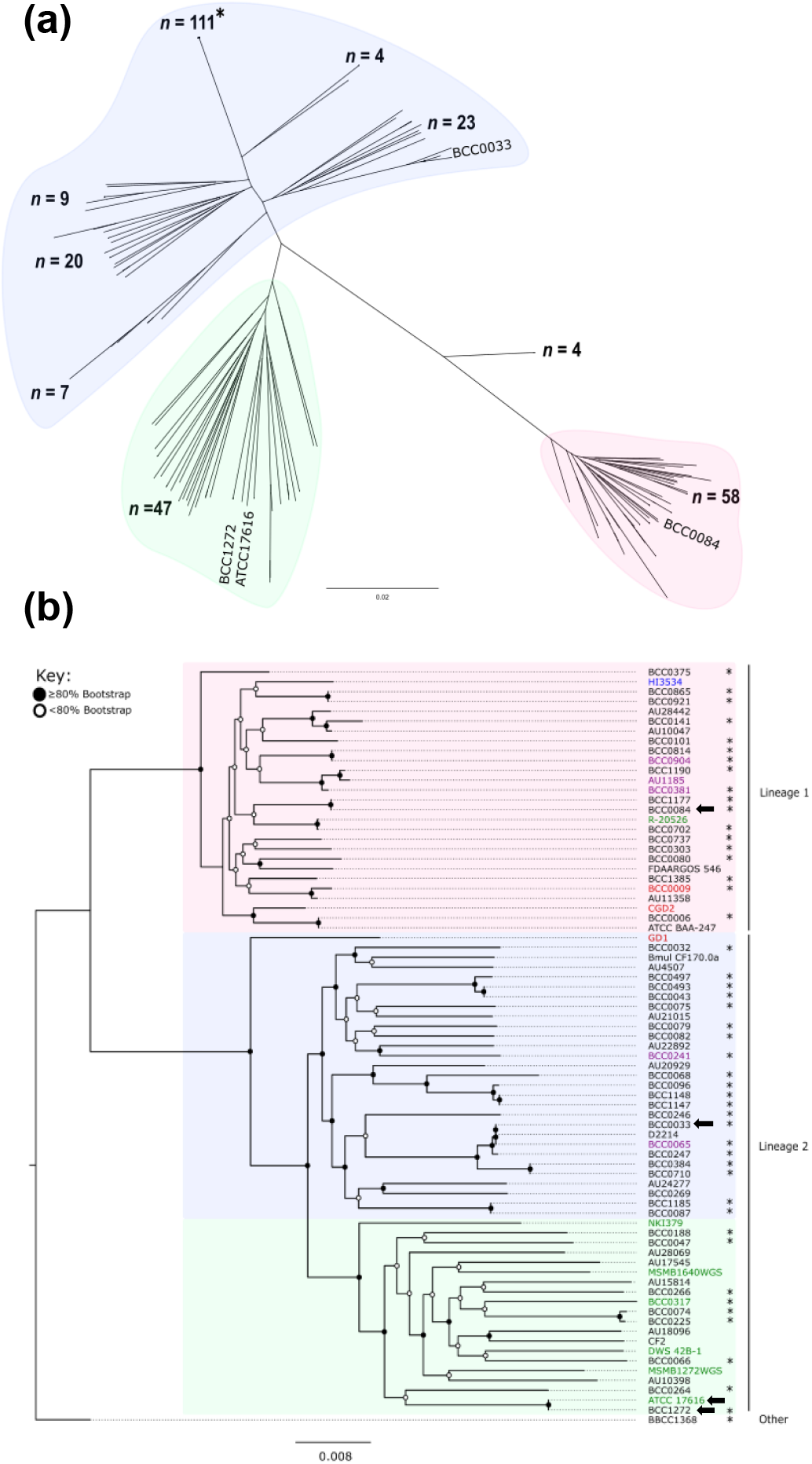
Core gene phylogenetic analysis of *B. multivorans* genomes corroborates the presence of two major lineages. **(a)** A core-gene phylogeny of 283 *B. multivorans* strains was generated by aligning 4319 core genes using RAxML (100 bootstraps). The tree was rooted using BCC1368 (black arrow) and comprised an outgroup of 4 isolate genomes. Lineage 1 (red), 2a (green) and 2b (blue) groups are shaded. The position of the selected model *B. multivorans* strains is indicated by the strain names. The single strain group (*n =* 111) represents sequential isolates of CF strain sequenced during chronic *B. multivorans* infection. (b) The core gene phylogeny of the 77-strain panel is also presented (also an alignment of the 4,319 core-genes using RAxML with 100 bootsraps). Nodes have been allocated a white circle to illustrate ≤80% bootstrap or a black circle for ≥80% bootstrap. The lineages are labelled (right) and isolates indicated with the asterisk were genome sequenced and studied phenotypically as part of this study (see Table 1). Isolate strain names are provided and the text colour denotes their source (Black = CF, green = ENV, blue = ENVH, purple = NON-CF, and red = CGD); the position of the model strains is indicated by the black arrows. The number of base substitutions per site are indicated by the scale bars on each respective phylogeny.

**Figure 3.**
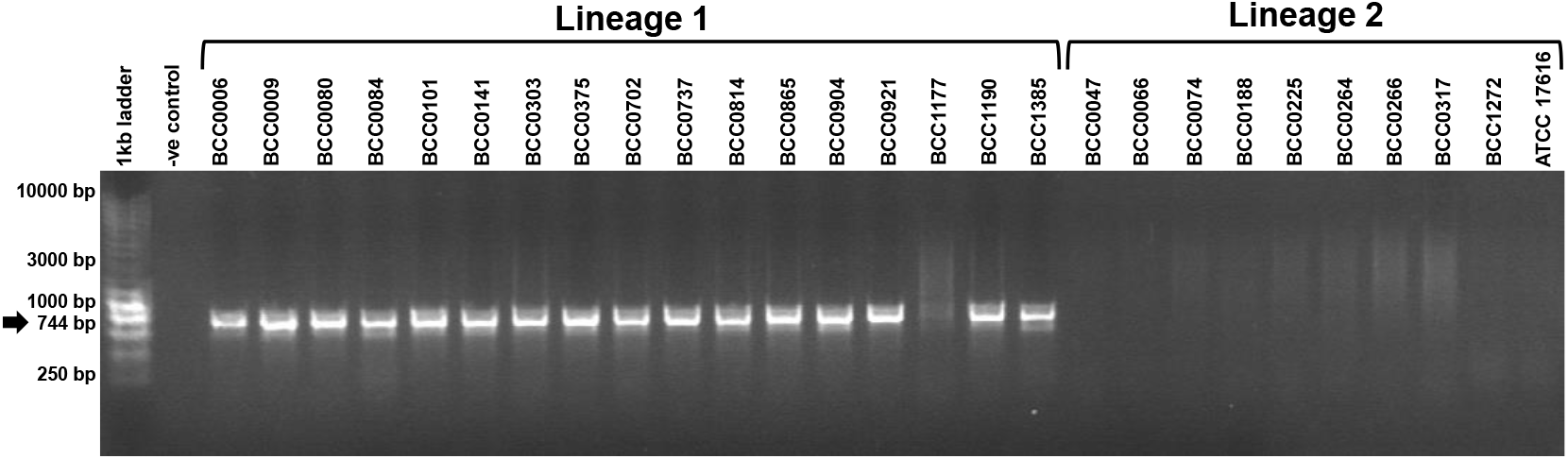
Specificity of the *ghrB_1* for identification of lineage 1 *B. multivorans* strains. The correct PCR amplicons (744 bp; see arrow on right) resulting from a *ghrB_1* PCR on 18 lineage *B. multivorans* strains is shown (strain names are shown above each lane). No amplicon products were produced from the *B. multivorans* lineage 2 strains (10 shown on the gel) or the water negative control. Molecular size ladders (1 kb ladder) are shown with the relevant size DNA fragments labelled. The BCC1177 DNA sample in this specific PCR was slightly degraded but worked when repeated by PCR.

The selected sub-panel of 77 *B. multivorans* strains demonstrated the same phylogenomic population biology and 2 lineage split (Figure 2b). The greater diversity within lineage 2 strains was characterised by the longer branch length compared to lineage 1 strains, with the split into 2a and 2b sub-groups clearly observed (Figure 2b). Interestingly, although the total number of environmental isolates of *B. multivorans* was low in the larger 283 genome dataset (*n* = 23; including *n* = 2 ENVH strains), a total of 20 environmental isolates clustered within lineage 2 (16 within the 2a subgroup and 4 in 2b; Table S1). The localisation of six of these ENV lineage 2 genomes, and one lineage 1 ENV strain is shown in the core gene sub-panel phylogeny (Figure 2b). These data corroborate previous findings that *B. multivorans* is a Bcc species that is rarely isolated from the natural environment [8].

Multlilocus sequencing typing has been a key epidemiological resource from which to understand *Burkholderia* infection on a global scale [55], with the Bcc MLST [23] database currently comprising over 4000 *B. multivorans* isolate profiles. Therefore, the phylogenomic divisions based on 4319 core genes were evaluated against the 7-gene phylogenies from Bcc MLST strain typing scheme [23]. The MLSTcheck program [40] was implemented to derive an MLST allele profile and ST for the strain panel genomes (Table 2). Within the newly sequenced strains, this revealed four novel alleles (BCC0082 [2 alleles], BCC0266, and BCC0737) and four novel STs, with a total of 43 unique STs within the 77-strain panel (Table 2). There were six different clonal complexes (CC) observed within the strain panel, with six strains part of CC1. This CC encompassed ST15 and ST16 *B. multivorans* strains which had caused outbreaks of CF infection in several countries [22]. While phylogenetic analysis of the seven concatenated MLST alleles was able to resolve a two-lineage split within *B. multivorans*, a subset of strains clustered differently and flipped between the 2a and 2b subgroups (Figure S1) that had been assigned by the core gene analysis (Figure 2a). This demonstrated that the limited resolution of MLST would not be able to accurately cluster within lineage 2 strains but could assign them to the overall group. It also confirmed that recombination observed within the seven MLST loci [22] is a feature of *B. multivorans*.

### Design and testing of *B. multivorans* lineage-specific PCRs

To enable rapid identification and future epidemiological surveillance of the *B. multivorans* lineages, PCR diagnostics were designed and evaluated as follows. Following a pan-GWAS analysis [49], three genes were identified as 100% present and specific to lineage 1 strains: *yiaJ_1*, a predicted DNA-binding transcriptional repressor, *ghrB_1*, a putative glyoxylate/hydroxypyruvate reductase B, and *naiP_3*, a predicted niacin/nicotinamide transporter (Table S3). All three genes were encoded on the second chromosomal replicon when compared to the complete genome of the lineage 2 CF strain *B. multivorans* BCC0084. A single target gene, *glnM_2*, a putative glutamine ABC transporter permease, was specific to lineage 2 *B. multivorans* genomes and encoded on replicon 1, when correlated to the complete genome of strain ATCC 1716 (Table S3). After BLAST analysis of *in silico* primer specificity and consideration of mismatches in the primer designs (Table S3), the *ghrB_1* and *glnM_2* PCRs (Table 3) were tested against the panel of 49 phenotypically analysed strains (Table 1). Each PCR demonstrated specificity, with the correct amplicon size produced for strains of the target lineage, and they did not amplify the opposing *B. multivorans* lineage or control *B. ambifaria* and *B. cenocepacia* DNA (Figure 4; a *ghrB_1* PCR example).

**Figure 4.**
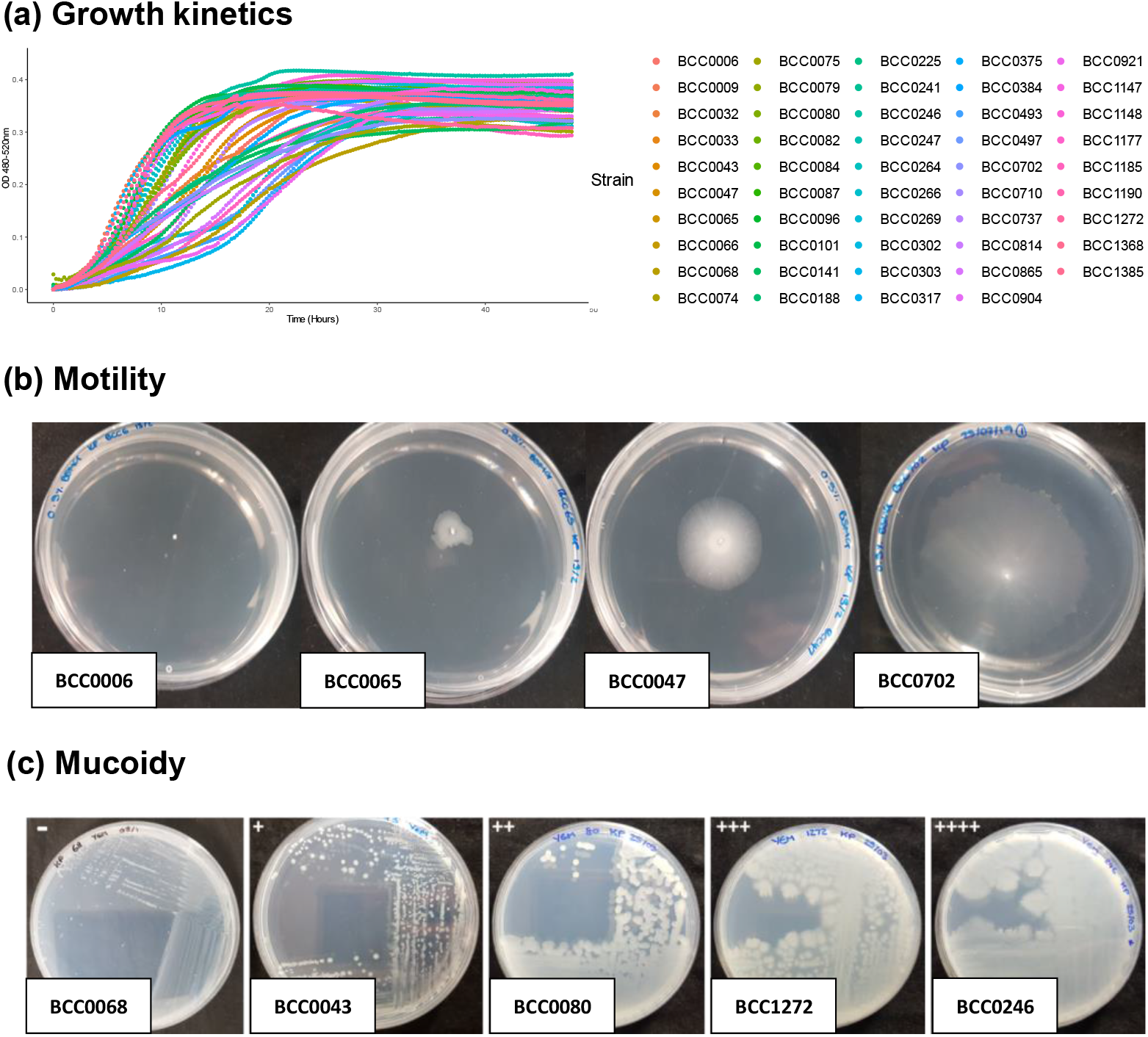
Phenotypic characteristic of *B. multivorans*. Panel A shows the growth curves measured using a Bioscreen C instrument for each of the 49 *B. multivorans* panel strains (and one additional strain). The mean optical density of technical (n=3) and biological replicates (n=3) is plotted for every 15 minute reading across 48 h. The key provides the strain names. Panel B shows the motility of selected *B. multivorans* strains ranging from low to high motility on 0.5% swarming BSM-G agar (BCC0006 = non-motile, BCC0065 = low motility, BCC0047 = intermediate motility, and BCC0702 = high motility). This panel represents the motility categories (diameters) which were observed on 0.3% swimming agar also. Panel C shows a section of *B. multivorans* strain reflective of the EPS production scale seen after growth on YEM agar.

### The *B. multivorans* phenotype is variable between strains and lineages

To examine the extent that the genomic lineages correlated to phenotypic differences *in vitro*, 49 representative strains (Table 1) were examined for growth kinetics, motility, biofilm formation, exopolysaccharide production and protease production. This collection comprised 18 lineage 1 strains, 30 lineage 2 strains (2a, n = 9; 2b, n = 21), and the outlier *B. multivorans* BCC1368. Analysis of growth kinetics demonstrated that all *B. multivorans* strains produced typical sigmoidal growth curves in TSB but varied in their growth characteristics (Figure 4a). In terms of maximum growth rate (collection mean = 0.032 h^-1^), 11 strains (BCC0032, BCC0068, BCC0075, BCC0188, BCC0225, BCC0247, BCC0375, BCC00497, BCC0702, BCC0814 and BCC0865; 22%) fell below the first quartile were designated as slow growing (Table S4). Outliers for lag phase (collection mean = 5.02 h) were BCC0303, BCC0269, BCC1185, BCC0493 and BCC0921 (mean = 11.16 h) which possessed prolonged lag phases and small colony phenotypes on TSA (except for BCC0269) (Table S4).

Motility on nutrient (TSA) versus minimal medium (BSM-G) was examined for swimming and swarming phenotype. A consistent finding was that the majority of *B. multivorans* strains were motile on at least one type of agar (96%; 47 of 49; Figure 4b), but BCC0068 (a CF isolate) and BCC0904 (a non-CF infection isolate) were non-motile on all agar types (Table S5). Overall, a greater number of *B. multivorans* strains had the ability to swim (87%) rather than swarm (80%) on at least one medium type (Table S5). No statistically supported phenotypic differences were found between lineages in relation to motility (Figure S2). The majority of *B. multivorans* strains (42 of 49; 86%) were able to form biofilms *in vitro* within the 96-well PVC-plate binding assay. A previous study [56] had shown strain ATCC 17616 to be a high biofilm former and BCC0010 (also known as strain C1962) to be a weak biofilm former. Three strains formed more biofilm than ATCC 17616 (BCC0047, BCC1147 and BCC1272), while 7 *B. multivorans* strains had an average biofilm formation less than BCC0010 (BCC0068, BCC0075, BCC0264, BCC0493, BCC0814, BCC0865 and BCC0921). The ability to form biofilms *in vitro* was not statistically linked to each lineage (Figure S3).

Using the semi-quantitative YEM agar assay to determined exopolysaccharide production [45], the majority of *B. multivorans* tested (79 of 84; including all the 49 panel strains in Table 1) had the ability to produce mucoid phenotypes on YEM agar (Figure 4b). The non-mucoid phenotype was only observed within five strains (BCC0006, BCC0068, BCC0188, BCC0493 and BCC0497), and interestingly, four of these strains also exhibited no or low motility on all agars (Table S5). All 49 *B. multivorans* (Table 1) strains were assessed for protease production using an updated assay [46], but none were found to secrete active proteases *in vitro*. In contrast, the positive control, *P. aeruginosa* strain LES B58, produced a clear halo of protease activity on all assays.

### Selection of *B. multivorans* model CF strains

Using the resource of extensive phylogenomic and phenotypic analyses obtained, three model *B. multivorans* CF strains were selected. The criteria used accounted for phylogenomic lineage and the possession of a phenotype reflective of the majority of *B. multivorans* strains. The selected strains were: BCC0033 (also known as C5568) as a lineage 1 CF strain from Canada that was representative of the globally spread ST-16 and clonal complex 1 (Table 2); BCC0084 (also known as C6398), a lineage 2b CF strain from Canada (ST-195; Table 2), and BCC1272 (also known as AU0453), a lineage 2a CF strain from the USA (ST-21; Table 2). In addition to these three CF strains, the *B. multivorans* reference strain ATCC 17616 (BCC0011), a lineage 2a soil isolate was considered as a fourth model strain because of its well-studied nature. Although ATCC 17616 was isolated from soil, CF isolate BCC1272 had the same MLST type, ST-21. Core-gene phylogenomic analysis (Figure 2b), and complete sequence analysis (Table 1) also showed that the soil and CF isolate were essentially identical at genomic level. All the four model strains were also shown to be genetically amenable to plasmid transformation by successful electroporation and reporter gene expression from plasmid pIN301-eGFP and pIN233-mCherry [47]. Finally, to ensure genomic resources for the model CF strains BCC033, BCC0084 and BCC1272 were substantive, they were subjected to complete genome sequencing (see Table 1 for accession numbers).

### The *B. multivorans* model strains were capable of murine respiratory infection

To understand if the selected model *B. multivorans* were proficient in their ability to colonise the mammalian nasopharynx and lung, and therefore suitable for pathogenicity/therapeutic testing, they were examined in a murine model of respiratory infection [19, 20, 51]. A single experiment with statistical power to evaluate basic bacterial survival kinetics was carried out using strains BCC0033 and its eGFP::pIN301 derivative (BCC033-GFP), BCC0084 eGFP::pIN301 (BCC084-GFP) and *B. multivorans* ATCC 17616. All the initial *B. multivorans* stocks used for infection and the pooled isolates obtained from 3- and 5-days post infection (the nasopharynx and lungs), were subjected to Illumina re-sequencing to confirm their genetic identity and evaluate if short-term genomic evolution had occurred.

Intranasal infection with approximately 10^7^ c.f.u. ml^-1^ of each *B. multivorans* strain resulted in colonisation of the respiratory tract ranging from 10^2^ to 10^5^ log_10_ c.f.u. within both the nasopharynx and lungs, which persisted over the 5-day infection (Figure 5). In rank order, BCC0084-GFP had the greatest rate of lung colonisation (1.8 × 10^4^ to 1.7 × 10^5^ c.f.u. ml^-1^) over 5 days, followed by BCC0033 (1.3 × 10^4^ to 2.5 × 10^4^ c.f.u. ml^-1^), BCC0033-GFP (7.9 × 10^3^ to 1.5 × 10^4^ c.f.u. ml^-1^) and strain ATCC 17616 which possessed the lowest lung infection rate (1.1 × 10^2^ to 1.8 × 10^3^ c.f.u. ml^-1^) (Figure 5).

**Figure 5.**
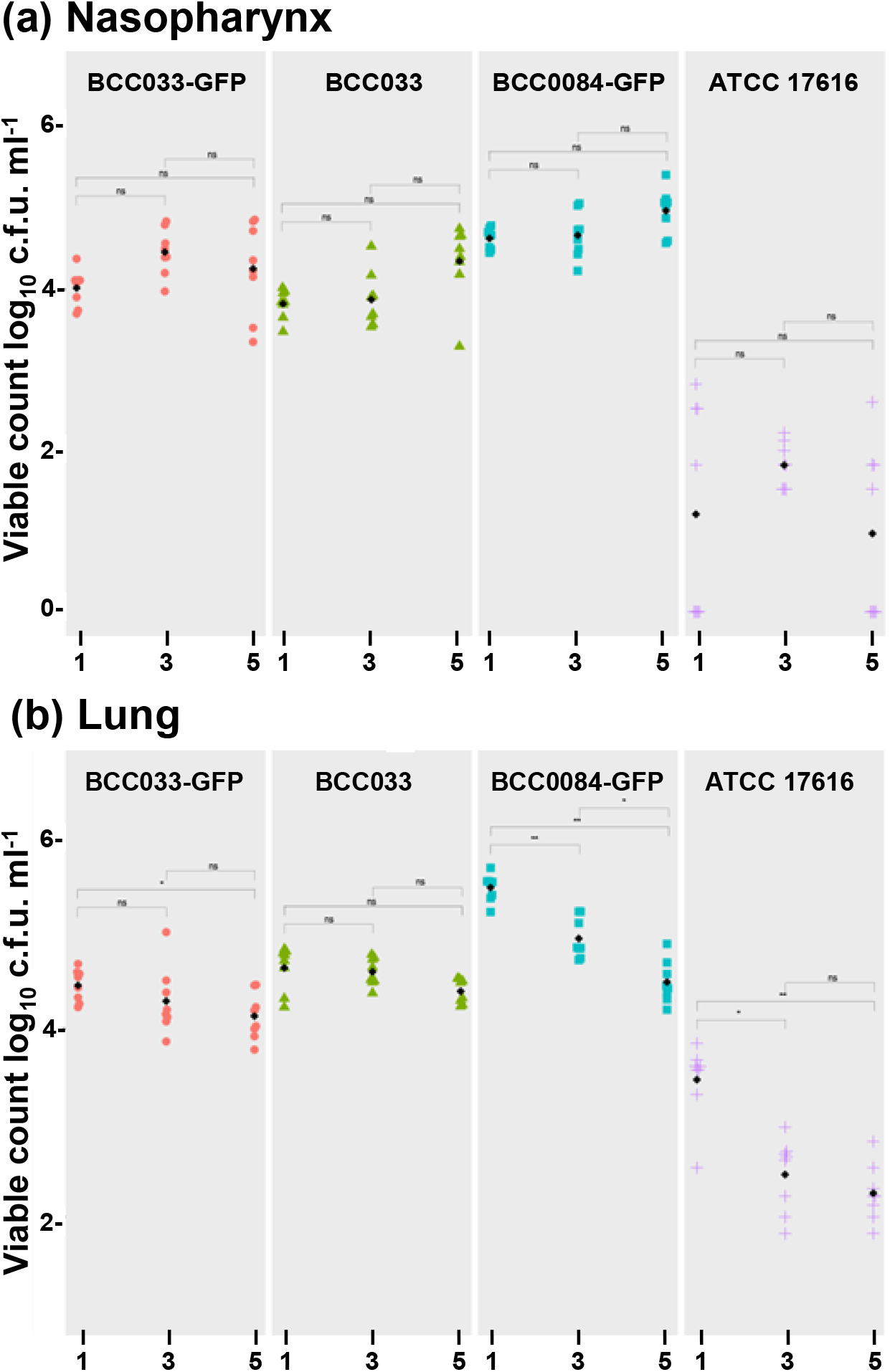
*Burkholderia multivorans* model strains can persist within a mammalian respiratory infection model. Mouse lung and nasopharynx infection dynamics for the selected *B. multivorans* model strains are shown. Viable counts (c.f.u.) for the *B. multivorans* strains at days 1, 3 and 5 post-infections are shown with the within strain statistical significance indicated for each time point. The panels show: (A) infection of the nasopharynx and (B) the lungs, with the individual (coloured) and median (black) c.f.u. for each tissue plotted.

Genome resequencing of the pooled isolates from the nasopharynx and lung demonstrated that infection isolates were essentially isogenic with each respective inoculated strain. Scaffolding of the short-read sequences to the complete genomes demonstrated that no major genomic rearrangements had occurred during the short-term infection. Overall, 242 SNP variants were observed to have accumulated amongst the four *B. multivorans* genomes as follows. In total, 72 (29.75%) had annotated effects that were: 4 conservative in-frame insertions, 4 disruptive in-frame deletions, 26 missense variants, 4 stop lost and splice region variants and 34 synonymous mutations. *B. multivorans* ATCC 17616 harboured the greatest number of SNPs (n = 110). A total of 27 and 20 SNPs were found in the pooled lung isolates at day 3 and 5, respectively, and 23 and 40 SNPs in the nasopharynx, at the same respective time points. Strain BCC0033 harboured the fewest SNPs with 27 identified (four in lung day 3, seven in lung day 5, eight in the nasopharynx on days 3 and 5), followed by BCC0033-GFP with 33 SNPs with a similar distribution (seven SNPs in lung day 3, fourteen SNPs in lung day 5 and six SNPs in both nasopharynx days 3 and 5). BCC0084 eGFP::pIN301 had a total of 72 SNPs (nineteen in lung day 3, seventeen in lung day 5, sixteen in nasopharynx day 3, and twenty in nasopharynx day 5).

## Discussion

A limited number of *Burkholderia* species have been subjected to in-depth population biology, phylogenomic and phenotypic analysis. *B. multivorans* has been previously investigated to MLST level demonstrating the presence of globally distributed clonal complexes [22]. Using genomic analyses, we have taken epidemiological understanding a step further, identifying two evolutionary lineages within *B. multivorans*. Although no difference in the distribution of CF isolates across the two *B. multivorans* lineages was seen, it is interesting that the majority of globally distributed *B. multivorans* clonal complexes [22] resided in lineage 2b (Table 2). In comparison to *B. cenocepacia* [9], there are no currently defined model CF strains for *B. multivorans*. By combining the genomic findings with the common phenotypic features of *B. multivorans*, three model CF strains were identified as suitable for future studies alongside the well-characterised soil isolate ATCC 17616. The model strains (BCC0033, BCC0084 and ATCC17616) were all capable of *in vivo* infection in a murine model of respiratory tract infection, providing a future platform for virulence analysis and therapeutic screening. With straightforward PCR diagnostic probes also designed to rapidly identify each *B. multivorans* genomic lineage, clinical laboratories now have straightforward tools to evaluate their associated epidemiology.

Several *B. cepacia* complex species have recently been observed to contain unexpected genomic diversity, resulting in the identification of novel genomic taxa within them. For example, the historical *recA* gene-based lineage originally identified in *B. cenocepacia* as III-B [11], was identified as a separate genomic taxa [12] and subsequently proposed as the new species *B. orbicola* sp. nov. [13]. *B. gladioli*, the third most common *Burkholderia* CF pathogen seen in the US [6] was thought to comprise several pathovars, but genomic analyses demonstrated that five distinct evolutionary clades existed within this single genomic species [21]. Further, bongkrekic acid toxin producing strains (clades 1a, 1b and 1c) occurring as CF lung infections were identified for the first time within *B. gladioli*. Finally, across *Burkholderia* species as a whole, multiple novel genomic taxa have been identified, with only approximately half of these having formal species names [36]. Our phylogenomic analysis of *B. mutlivorans* shows that this important CF pathogen does not harbour further genomic taxa (Figure 1), but does comprise 2 major evolutionary lineages (Figure 2). Like the two genomic groups observed in the major CF pathogen *P. aeruginosa* [57, 58], the pathogenic significance of these *B. multivorans* lineages remains to be determined.

We identified that *B. multivorans* strains possess highly variable phenotypes, with no direct linkage to their genomic lineage. However, what was consistent was that most strains from CF infection were motile, able to form biofilms *in vitro*, but lacked the ability to produce proteases on growth media. When investigating the *B. multivorans* growth rate *in vitro*, two strain groups were apparent, splitting the isolates into approximately two groups, those that reached stationary phase by 24 h, versus those reaching this growth stage at 30 h (Figure 4a). Reduced *B. multivorans* growth rates have previously been observed in CF infection [16] and is also the case for *P. aeruginosa* chronic lung infection isolates [59]. All 11 *B. multivorans* strains identified as slow growers had been recovered from CF infection, suggesting this is also pathogenic adaptation the species makes during chronic infection.

Overall, screening a collection of *B. multivorans* demonstrated that the majority of strains retained motility as a core phenotype. This contrasts with *P. aeruginosa*, where isolates from chronic CF lung infection are known to become non-motile [60], but correlates with longitudinal analysis of *B. cepacia* complex isolates, where just swimming motility was examined [61]. Non-swimming *B. multivorans* were rare among the collection of isolates screened (14%) and loss of swimming motility was previously suggested as not a common adaptive feature of chronically infecting CF strains [61]. Silva, Santos [16] examined 22 longitudinal isolates recovered from an individual with CF spanning 20 years and showed decreased swimming motility of this single strain that was likely due to mutations accumulating in the cyclic di-GMP (c-di-GMP) metabolism pathway. Loss of motility has been observed in invasive *B. cenocepacia* strains that were isolated from the bloodstream of CF individuals suffering with acute ‘cepacia syndrome’ [62]. Of the genetically diverse isolates screened in our study, only *B. multivorans* strain BCC0068 (a CF isolate) was non-motile on all motility agar types, while BCC0006 showed no swarming motility (Figure 4b), but retained limited swimming ability (Table S5). For *B. cepacia* complex species, it has been shown that infection with nonmucoid strains correlates to an increased lung function decline, as compared to infection with mucoidal variants [14]. Only five nonmucoid *B. multivorans* variants were identified in our study, but all the nonmucoid strains exhibited no or limited motility, as had been observed in other studies [15, 61, 63].

A useful finding from the *B. multivorans* strains examined using the murine respiratory infection model was that they demonstrated initial levels of lung and nasopharynx colonisation similar to *P. aeruginosa* strain LESB65 (between log 2 and 4 c.f.u. in each tissue) [19, 51]. This is substantially greater than the low level of colonization (<1000 c.f.u/tissue) observed for the *B. cepacia* complex species, *B. ambifaria*, in the same murine infection model [20]. The limited ability of *B. ambifaria* to colonise the mammalian respiratory tract correlates to the species epidemiology in CF, where it has historically been rarely seen [6] or more recently, not observed [5] compared to *B. multivorans*, which was the dominant CF *Burkholderia* in both epidemiological studies. The *B. multivorans* CF strain BCC0084-GFP (lineage 1) was the most adept coloniser of both the mouse lung and nasopharynx, with the environmentally derived ATCC 17616 showing the lowest colonisation rate (Figure 5); however, this was still greater, in terms of infectivity, compared to *B. ambifaria* [20]. Further systematic studies of *B. cepacia* complex species in this murine model of infection [19, 20, 51] will need to be carried out to establish their comparative pathogenicity, but promisingly, it is clear the model is a good system for studying *B. multivorans*.

In summary, although *B. multivorans* possesses a highly variable phenotype, it is genomically one species harbouring 2 major lineages. At this stage in our analyses, no differences between *B. multivorans* lineages have been observed. However, with the identification of representative model strains reflecting each lineage and the conserved species phenotypes, as well as PCR primers to rapidly identify each lineage, in depth studies of *B. multivorans* as a CF can now be undertaken.

## Supporting information

Supplementary date file

## Acknowledgements

KMP was funded by PhD studentship award from the Medical Research Council GW4 Biomed Doctoral Training Program (Project No. BV19101122). EM acknowledges additional CF lung infection research support from the US Cystic Fibrosis Foundation (grant MAHENT20G0). AEG and DRN were supported by a Sir Henry Dale fellowship, awarded to DRN and funded by Wellcome and the Royal Society (204457/Z/16/Z). EM and DRN note support of their research from a UK Strategic Research Centre (SRC) “An evidence-based preclinical framework for the development of antimicrobial therapeutics in cystic fibrosis” (PIPE-CF; Project No. SRC 022) funded by UK Cystic Fibrosis Trust and US Cystic Fibrosis Foundation. TRC Acknowledges support from the MRC for the CLIMB-BIG-DATA resource (Grant Reference: MR/T030062/1).

## Author contributions

The CReDIT contributor roles taxonomy was used to recognise author contributions as follows:

Conceptualization: EM, TRC, KMP

Methodology: KMP, EM, AEG, DRN

Software: KMP, TRC

Validation: KMP, EM, AEG, DRN

Formal analysis: KMP, AEG, DRN

Investigation: KMP, EM, AEG, DRN

Resources: EM, TRC, DRN

Data Curation: KMP

Writing – original draft preparation: KMP, EM

Writing – review and editing: all authors

Visualization: KMP, EM

Supervision: EM, DRN

Project administration: EM, DRN

Funding acquisition: EM, TRC, DRN, KMP

