## Supplementary date file for "Identification of two distinct phylogenomic lineages and model strains for the understudied cystic fibrosis lung pathogen *Burkholderia multivorans*"

**Affiliations:**

**Current affiliations:**

[3] Big Data Institute, Nuffield Department of Population Health, Li Ka Shing Centre for Health Information and Discovery, Old Road Campus, University of Oxford, Oxford OX3 7LF, UK

### Supplementary tables

Table S1. Details of the 283 *B. multivorans* genome sequences analysed in this study

Table S2 Quality statistics of the 73 *B. multivorans* draft bacterial genomes sequenced in this study

Table S3. PCR primer sequences designed for all four *B. multivorans* lineage-specific target genes

Table S4. Growth kinetics of the *B. multivorans* strains showing growth rate, lag phase and maximum OD

Table S5. Swimming and swarming motility within the *B. multivorans* strains

Table S6. Colony morphology and EPS production of *B. multivorans*

### Supplementary Figures

Figure S1. MLST-gene phylogeny of the 77 genome *B. multivorans* strain panel

Figure S2. Swimming and swarming motility in the B. multivorans strain panel after 24 h incubation at 37$^{o}$C

Figure S3. Biofilm formation of the *B. multivorans* strain panel (n = 49) after 24 h

### Table S1. Details of the 283 *B. multivorans* genome sequences analysed in this study

| Strain | Isolate Type | Lineage | Accession Number | Sequence Type (ST) | Other name | Origin |
|---|---|---|---|---|---|---|
| 701_BMUL | NA | 2b | GCF_001058025.1 | 16 | | Washington, USA |
| 800_BMUL | NA | 2b | GCF_001058485.1 | 16 | | Washington, USA |
| ATCC_17616 | ENV | 2a | GCF_000018505.1 | 21 | | USA |
| ATCC_BAA-247 | CF | 1 | GCF_000959525.1 | 650 | | Brussels, Belgium |
| AU10047 | CF | 1 | GCF_002980655.1 | | | USA |
| AU10086 | CF | 2b | GCF_002980675.1 | 15 | | USA |
| AU10398 | CF | 2a | GCF_002980695.1 | | | USA |
| AU10897 | CF | 2b | GCF_002980715.1 | | | USA |
| AU11204 | CF | 1 | GCA_002980785.1 | | | USA |
| AU11233 | CF | 2a | GCF_002980775.1 | | | USA |
| AU11358 | CF | 1 | GCF_002981015.1 | | | USA |
| AU11772 | CF | Other | GCF_002981035.1 | | | USA |
| AU1185 | NON | 1 | GCF_001718755.1 | 18 | | USA |
| AU12481 | CF | 2a | GCF_002981075.1 | | | USA |
| AU13919 | CF | 2b | GCF_002981095.1 | | | USA |
| AU14328 | CF | 2b | GCF_002980825.1 | | | USA |
| AU14364 | CF | 2a | GCF_002980815.1 | | | USA |
| AU14371 | CF | 2b | GCF_002980855.1 | | | USA |
| AU14786 | CF | Other | GCF_002980875.1 | | | USA |
| AU15814 | CF | 2a | GCF_002980895.1 | | | USA |
| AU15954 | CF | 2b | GCF_002980905.1 | 190 | | USA |
| AU16734 | CF | 1 | GCF_002981615.1 | | | USA |
| AU17135 | CF | 1 | GCF_002981135.1 | | | USA |
| AU17534 | CF | 2a | GCF_002980935.1 | | | USA |
| AU17545 | CF | 2a | GCF_002980995.1 | | | USA |
| AU18096 | CF | 2a | GCF_002981145.1 | | | USA |
| AU19518 | CF | 2a | GCF_002981195.1 | | | USA |
| AU19564 | CF | 2b | GCF_002981155.1 | | | USA |
| AU19659 | CF | 1 | GCF_002981255.1 | | | USA |
| AU19729 | CF | 2a | GCF_002981215.1 | | | USA |
| AU20929 | CF | 2b | GCF_002981635.1 | | | USA |
| AU21015 | CF | 2b | GCF_003048355.1 | | | USA |
| AU21596 | CF | 2b | GCF_002981675.1 | | | USA |
| AU21747 | CF | 2a | GCF_002981315.1 | | | USA |
| AU22436 | CF | 1 | GCF_002981695.1 | | | USA |
| AU22892 | CF | 2b | GCF_002981295.1 | 190 | | USA |
| AU23365 | CF | 1 | GCF_002981715.1 | | | USA |
| AU23668 | CF | 2a | GCF_002981725.1 | | | USA |
| AU23690 | CF | 1 | GCF_002981325.1 | | | USA |
| AU23919 | CF | Other | GCF_002981335.1 | | | USA |
| AU23995 | CF | 2b | GCF_002981755.1 | | | USA |
| AU24277 | CF | 2b | GCF_002981375.1 | | | USA |
| AU25057 | CF | 1 | GCF_002981775.1 | | | USA |
| AU25543 | CF | 2a | GCF_002981815.1 | | | USA |
| AU26250 | CF | 2b | GCF_002981835.1 | 190 | | USA |
| AU27706 | CF | 1 | GCF_002981395.1 | 190 | | USA |
| AU28069 | CF | 2a | GCF_002981845.1 | | | USA |
| AU28442 | CF | 1 | GCF_002981415.1 | | | USA |
| AU29198 | CF | 2b | GCF_002981455.1 | | | USA |
| AU30050 | CF | 1 | GCF_002981485.1 | | | USA |
| AU30438 | CF | 1 | GCF_002981515.1 | | | USA |
| AU30441 | CF | 1 | GCF_002981535.1 | | | USA |
| AU30760 | CF | 2b | GCF_002981555.1 | | | USA |
| AU4507 | CF | 2b | GCF_002981595.1 | | | USA |
| BCC0005 | CF | 2b | ERS785011 | 19 | LMG 18822; C5393 | British Colombia, Canada |
| BCC0006 | CF | 1 | ERZ1645180 | 270 | 4F | Toronto, Canada |

| | | | | | | |
|---|---|---|---|---|---|---|
| BCC0008 | CF | 2b | ERS784904 | 27 | LMG 16660; C1576 | Glasgow, UK |
| BCC0009 | CGD | 1 | ERZ1645179 | ND | LMG 18824 | USA |
| BCC0010 | NON | 1 | ERS784919 | 18 | LMG 16665; C1962; FC0762 | UK |
| BCC0031 | CF | 2b | ERS785002 | 191 | KE1; CO514 | British Colombia, Canada |
| BCC0032 | CF | 2a | ERZ1645178 | ND | C3865; HA1 | British Colombia, Canada |
| BCC0033 | CF | 2b | ERZ1645177 | ND | C5568 | California, USA |
| BCC0037 | CGD | 2a | ERS785053 | 271 | CEP0178 | California, USA |
| BCC0043 | CF | 2b | ERZ1645176 | ND | C1528 | Belfast, UK |
| BCC0047 | CF | 2a | ERZ1645175 | ND | C1712 | Aberdeen, UK |
| BCC0050 | ENV | 2b | ERS785036 | 493 | J0365 | UK |
| BCC0059 | CF | 2b | ERS785022 | 287 | CEP0494 | Montreal, Canada |
| BCC0065 | NON | 2b | ERZ1645174 | 16 | CEP0600 | Oregon, USA |
| BCC0066 | CF | 2a | ERZ1645173 | 24 | CEP0602 | British Colombia, Canada |
| BCC0067 | CF | 1 | ERS784935 | 189 | CEP0603 | Montreal, Canada |
| BCC0068 | CF | 2b | ERZ1645172 | 329 | CEP0604 | Montreal, Canada |
| BCC0074 | CF | 2a | ERZ1645170 | ND | HI-2308 | USA |
| BCC0075 | CF | 2b | ERZ1645169 | ND | C1579 | Glasgow,UK |
| BCC0079 | CF | 2b | ERZ1645168 | ND | C3430 | British Colombia, Canada |
| BCC0080 | CF | 1 | ERZ1645167 | ND | C4861 | British Colombia, Canada |
| BCC0082 | CF | 2b | ERZ1645166 | ND | C6100 | British Colombia, Canada |
| BCC0084 | CF | 1 | ERZ1645165 | 195 | C6398 | British Colombia, Canada |
| BCC0087 | CF | 2b | ERZ1645164 | ND | C6935 | British Colombia, Canada |
| BCC0089 | CF | 2b | ERS785033 | 191 | C7274 | British Colombia, Canada |
| BCC0093 | CF | 1 | ERZ1756586 | 1023 | C7363 | British Colombia, Canada |
| BCC0096 | CF | 2b | ERZ1645163 | ND | C7062 | British Colombia, Canada |
| BCC0099 | CF | 2b | ERS785069 | 305 | C7510 | British Colombia, Canada |
| BCC0101 | CF | 1 | ERZ1645162 | 304 | C8298 | British Colombia, Canada |
| BCC0102 | CF | 2b | ERS784954 | 190 | C8467 | British Colombia, Canada |
| BCC0115 | CF | 1 | ERS784971 | 269 | Patient Q | Washington, USA |
| BCC0134 | CF | 2a | ERZ1756605 | 1088 | CEP0699 | Denmark |
| BCC0141 | CF | 1 | ERZ1645161 | 198 | CEP0740; AU0071 | USA |
| BCC0149 | CF | 1 | ERS785050 | 17 | FC0328 | USA |
| BCC0175 | CGD | 1 | ERS785024 | 355 | FC0442 | Colorado, USA |
| BCC0181 | CF | 2b | ERZ1756585 | 16 | LMG 14273 | Belgium |
| BCC0188 | CF | 2a | ERZ1645171 | 196 | LMG 14276 | Belgium |
| BCC0225 | CF | 2a | ERZ1645159 | ND | CEP0837 | Manitoba, Canada |
| BCC0241 | NON | 2b | ERZ1645160 | 272 | CEP0935 | Minnesota, USA |
| BCC0244 | CF | 2a | ERS785004 | 188 | CEP0938 | British Colombia, Canada |
| BCC0246 | CF | 2b | ERZ1645158 | 273 | NU9366 | New Zealand |

| | | | | | | |
|---|---|---|---|---|---|---|
| BCC0247 | CF | 2b | ERZ1645157 | 16 | CEP0949 | New Zealand |
| BCC0255 | CF | 2b | ERZ1756587 | ND | CEP0966 | Australia |
| BCC0264 | CF | 2a | ERZ1645156 | 274 | CEP0992 | New South Wales, Australia |
| BCC0266 | CF | 2a | ERZ1645155 | 354 | CEP0995 | New South Wales, Australia |
| BCC0269 | CF | 2b | ERZ1645154 | 275 | CEP1000 | New South Wales, Australia |
| BCC0292 | CF | 1 | ERZ1756588 | ND | C1409; CEP0133 | Edinburgh, UK |
| BCC0293 | CF | 1 | ERZ1756589 | 487 | C1406; BCC0293 | Oklahoma, USA |
| BCC0300 | CF | 2b | ERZ1756590 | 16 | STRAIN 88; CEP0503 | France |
| BCC0303 | CF | 1 | ERZ1645153 | 192 | AU0066 | USA |
| BCC0317 | ENV | 2a | ERZ1645152 | 25 | BP102 | British Colombia, Canada |
| BCC0321 | ENVH | 1 | ERZ1756591 | ND | C1664; CEP0181 | Manchester, UK |
| BCC0375 | CF | 1 | ERZ1645151 | 22 | AU0607 | USA |
| BCC0381 | NON | 1 | ERZ1645150 | 117 | LMG 16665 | UK |
| BCC0384 | CF | 2b | ERZ1645149 | 18 | FC0769 | British Colombia, Canada |
| BCC0470 | CF | 2b | ERZ1756592 | ND | MN (#303) | Italy |
| BCC0493 | CF | 2b | ERZ1645148 | 15 | BELF 1 (#353) | Belfast, UK |
| BCC0497 | CF | 2b | ERZ1645147 | 352 | BELF 5 (#357) | Belfast, UK |
| BCC0533 | CF | 2b | ERZ1756593 | 19 | C3168 | British Colombia, Canada |
| BCC0553 | CF | 2b | ERZ1756594 | 16 | C6558 | British Colombia, Canada |
| BCC0583 | CF | 2A | ERZ1756595 | 21 | C9140 | British Colombia, Canada |
| BCC0585 | CF | 2b | ERZ1756596 | 199 | C8814 | British Colombia, Canada |
| BCC0702 | CF | 1 | ERZ1645146 | 26 | IST453 | Portugal |
| BCC0704 | CF | 1 | ERZ1756597 | 836 | IST455 | Portugal |
| BCC0710 | CF | 2b | ERZ1645145 | 375 | A1-4 | Cardiff, UK |
| BCC0729 | CF | 2a | ERZ1756598 | 195 | C1628 | Cardiff, UK |
| BCC0737 | CF | 1 | ERZ1645144 | 15 | C1628 | UK |
| BCC0814 | CF | 1 | ERZ1645143 | 193 | 3-454 | Prague, Czech Republic |
| BCC0865 | CF | 1 | ERZ1645142 | 180 | 54-1138 | Prague, Czech Republic |
| BCC0901 | CF | 2b | ERZ1756599 | 182 | 90-N320 | Prague, Czech Republic |
| BCC0904 | NON | 1 | ERZ1645141 | 181 | 93-N331 | Prague, Czech Republic |
| BCC0907 | NON | 2a | ERZ1756600 | 630 | 96-N337 | Prague, Czech Republic |
| BCC0915 | CF | 1 | ERZ1756601 | 180 | 104-455 | Prague, Czech Republic |
| BCC0921 | CF | 1 | ERZ1645140 | 180 | 110-1226 | Prague, Czech Republic |
| BCC0962 | CF | 1 | ERZ1756602 | 180 | 151-454 | Prague, Czech Republic |
| BCC0968 | CF | 1 | ERZ1756603 | 181 | 157-1226A | Prague, Czech Republic |
| BCC1147 | CF | 2b | ERZ1645139 | 181 | C9861 | British Colombia, Canada |
| BCC1148 | CF | 2b | ERZ1645138 | 317 | C9862 | British Colombia, Canada |
| BCC1177 | CF | 1 | ERZ1645137 | 308 | D0913 | British Colombia, Canada |
| BCC1185 | CF | 2b | ERZ1645136 | 303 | D1348 | British Colombia, Canada |
| BCC1190 | CF | 1 | ERZ1645135 | 320 | D1442 | British Colombia, Canada |
| BCC1271 | CF | 2a | ERZ1756604 | ND | AU 4543 | USA |
| BCC1272 | CF | 2a | ERZ1645134 | 18 | HI 2229 | USA |
| BCC1367 | CF | 1 | ERZ1756606 | 25 | AU0623 | USA |
| BCC1368 | CF | Other | ERZ1645133 | 21 | AU1187 | USA |
| BCC1384 | CF | 1 | ERZ1756607 | 25 | AU0508 | USA |
| BCC1385 | CF | 1 | ERZ1645132 | 179 | AU4608 | USA |
| BCC1421 | CF | 2b | ERZ1756608 | 15 | C1576 | Glasgow, UK |
| CF170.0a | CF | 2b | | | | Toronto, Canada |

| | | | | |
|---|---|---|---|---|
| CF170.10a | CF | 2b | | Toronto, Canada |
| CF170.10b | CF | 2b | | Toronto, Canada |
| CF170.10c | CF | 2b | | Toronto, Canada |
| CF170.10d | CF | 2b | | Toronto, Canada |
| CF170.10e | CF | 2b | | Toronto, Canada |
| CF170.10f | CF | 2b | | Toronto, Canada |
| CF170.10g | CF | 2b | | Toronto, Canada |
| CF170.10h | CF | 2b | | Toronto, Canada |
| CF170.10i | CF | 2b | | Toronto, Canada |
| CF170.10j | CF | 2b | | Toronto, Canada |
| CF170.11a | CF | 2b | | Toronto, Canada |
| CF170.11b | CF | 2b | | Toronto, Canada |
| CF170.11c | CF | 2b | | Toronto, Canada |
| CF170.11d | CF | 2b | | Toronto, Canada |
| CF170.11e | CF | 2b | | Toronto, Canada |
| CF170.11f | CF | 2b | | Toronto, Canada |
| CF170.11g | CF | 2b | | Toronto, Canada |
| CF170.11h | CF | 2b | | Toronto, Canada |
| CF170.11i | CF | 2b | | Toronto, Canada |
| CF170.11j | CF | 2b | | Toronto, Canada |
| CF170.1a | CF | 2b | | Toronto, Canada |
| CF170.1b | CF | 2b | | Toronto, Canada |
| CF170.1c | CF | 2b | | Toronto, Canada |
| CF170.1d | CF | 2b | | Toronto, Canada |
| CF170.1e | CF | 2b | | Toronto, Canada |
| CF170.1f | CF | 2b | | Toronto, Canada |
| CF170.1g | CF | 2b | | Toronto, Canada |
| CF170.1h | CF | 2b | | Toronto, Canada |
| CF170.1i | CF | 2b | | Toronto, Canada |
| CF170.1j | CF | 2b | | Toronto, Canada |
| CF170.2a | CF | 2b | | Toronto, Canada |
| CF170.2b | CF | 2b | | Toronto, Canada |
| CF170.2c | CF | 2b | | Toronto, Canada |
| CF170.2d | CF | 2b | | Toronto, Canada |
| CF170.2e | CF | 2b | | Toronto, Canada |
| CF170.2f | CF | 2b | | Toronto, Canada |
| CF170.2g | CF | 2b | | Toronto, Canada |
| CF170.2h | CF | 2b | | Toronto, Canada |
| CF170.2i | CF | 2b | | Toronto, Canada |
| CF170.2j | CF | 2b | | Toronto, Canada |
| CF170.3a | CF | 2b | | Toronto, Canada |
| CF170.3b | CF | 2b | | Toronto, Canada |
| CF170.3c | CF | 2b | | Toronto, Canada |
| CF170.3d | CF | 2b | | Toronto, Canada |
| CF170.3e | CF | 2b | | Toronto, Canada |
| CF170.3f | CF | 2b | | Toronto, Canada |
| CF170.3g | CF | 2b | | Toronto, Canada |
| CF170.3h | CF | 2b | | Toronto, Canada |
| CF170.3i | CF | 2b | | Toronto, Canada |
| CF170.3j | CF | 2b | | Toronto, Canada |
| CF170.4a | CF | 2b | | Toronto, Canada |
| CF170.4b | CF | 2b | | Toronto, Canada |
| CF170.4c | CF | 2b | | Toronto, Canada |
| CF170.4d | CF | 2b | 1810 | Toronto, Canada |
| CF170.4e | CF | 2b | | Toronto, Canada |
| CF170.4f | CF | 2b | | Toronto, Canada |
| CF170.4g | CF | 2b | | Toronto, Canada |
| CF170.4h | CF | 2b | | Toronto, Canada |
| CF170.4i | CF | 2b | | Toronto, Canada |
| CF170.4j | CF | 2b | | Toronto, Canada |
| CF170.5a | CF | 2b | | Toronto, Canada |
| CF170.5b | CF | 2b | | Toronto, Canada |

| | | | | | |
|---|---|---|---|---|---|
| CF170.5c | CF | 2b | | | Toronto, Canada |
| CF170.5d | CF | 2b | | | Toronto, Canada |
| CF170.5e | CF | 2b | | | Toronto, Canada |
| CF170.5f | CF | 2b | | | Toronto, Canada |
| CF170.5g | CF | 2b | | | Toronto, Canada |
| CF170.5h | CF | 2b | | | Toronto, Canada |
| CF170.5i | CF | 2b | | | Toronto, Canada |
| CF170.5j | CF | 2b | | | Toronto, Canada |
| CF170.6a | CF | 2b | | | Toronto, Canada |
| CF170.6b | CF | 2b | | | Toronto, Canada |
| CF170.6c | CF | 2b | | | Toronto, Canada |
| CF170.6d | CF | 2b | | | Toronto, Canada |
| CF170.6e | CF | 2b | | | Toronto, Canada |
| CF170.6f | CF | 2b | | | Toronto, Canada |
| CF170.6g | CF | 2b | | | Toronto, Canada |
| CF170.6h | CF | 2b | | | Toronto, Canada |
| CF170.6i | CF | 2b | | | Toronto, Canada |
| CF170.6j | CF | 2b | | | Toronto, Canada |
| CF170.7a | CF | 2b | | | Toronto, Canada |
| CF170.7b | CF | 2b | | | Toronto, Canada |
| CF170.7c | CF | 2b | | | Toronto, Canada |
| CF170.7d | CF | 2b | | | Toronto, Canada |
| CF170.7e | CF | 2b | | | Toronto, Canada |
| CF170.7f | CF | 2b | | | Toronto, Canada |
| CF170.7g | CF | 2b | | | Toronto, Canada |
| CF170.7h | CF | 2b | | | Toronto, Canada |
| CF170.7i | CF | 2b | | | Toronto, Canada |
| CF170.7j | CF | 2b | | | Toronto, Canada |
| CF170.8a | CF | 2b | | | Toronto, Canada |
| CF170.8b | CF | 2b | | | Toronto, Canada |
| CF170.8c | CF | 2b | | | Toronto, Canada |
| CF170.8d | CF | 2b | | | Toronto, Canada |
| CF170.8e | CF | 2b | | | Toronto, Canada |
| CF170.8f | CF | 2b | | | Toronto, Canada |
| CF170.8g | CF | 2b | | | Toronto, Canada |
| CF170.8h | CF | 2b | | | Toronto, Canada |
| CF170.8i | CF | 2b | | | Toronto, Canada |
| CF170.8j | CF | 2b | | | Toronto, Canada |
| CF170.9a | CF | 2b | | | Toronto, Canada |
| CF170.9b | CF | 2b | | | Toronto, Canada |
| CF170.9c | CF | 2b | | | Toronto, Canada |
| CF170.9d | CF | 2b | | | Toronto, Canada |
| CF170.9e | CF | 2b | | | Toronto, Canada |
| CF170.9f | CF | 2b | | | Toronto, Canada |
| CF170.9g | CF | 2b | | | Toronto, Canada |
| CF170.9h | CF | 2b | | | Toronto, Canada |
| CF170.9i | CF | 2b | | | Toronto, Canada |
| CF170.9j | CF | 2b | | | Toronto, Canada |
| CF2 | CF | 2a | GCF_000286575.1 | 1079 | Unknown |
| CGD1 | CGD | 2b | GCF_000182255.1 | ND | Maryland, USA |
| CGD2 | CGD | 1 | GCF_000182275.1 | 442 | Maryland, USA |
| CGD2M | CGD | 1 | GCF_000182295.1 | 442 | Maryland, USA |
| D2095 | CF | 2b | GCF_000807825.1 | 16 | Vancouver, Canada |
| D2214 | CF | 2b | GCF_000807815.1 | 16 | Vancouver, Canada |
| DDS 15A-1 | ENV | 2a | GCF_000756005.1 | 802 | Australia |
| DSOPR54 | ENV | 2a | GCA_002222845.1 | | Singapore |
| DSOPR57 | ENV | 2a | GCA_002222875.1 | | Singapore |
| DWS_42B-1 | ENV | 2a | GCF_000756965.1 | 809 | Unknown |
| FDAARGOS_246 | CF | 1 | GCF_003019965.1 | | Unknown |
| FDAARGOS_546 | CLIN | 1 | GCF_003938705.1 | | Unknown |
| FDAARGOS_547 | CLIN | 2a | GCF_003812365.1 | | USA |
| FDAARGOS_548 | CLIN | 2a | GCF_003812585.1 | | Unknown |

| | | | | | | |
|---|---|---|---|---|---|---|
| HI3534 | ENVH | 1 | GCF_001528605.1 | 620 | | USA |
| MSMB1128WGS | ENV | 2b | GCF_001529795.1 | ND | | Australia |
| MSMB1272WGS | ENV | 2a | GCF_001529925.1 | 1088 | | Australia |
| MSMB1535WGS | ENV | 2b | GCF_001530485.1 | ND | | Australia |
| MSMB1640WGS | ENV | 2a | GCF_001718995.1 | 802 | | Australia |
| MSMB1641WGS | ENV | 2a | GCF_001531425.1 | 802 | | Australia |
| MSMB1794WGS | ENV | 2b | GCF_001526715.1 | ND | | Australia |
| MSMB1916WGS | ENV | 2a | GCF_001530985.1 | ND | | Australia |
| MSMB2008WGS | ENV | 2a | GCF_001528045.1 | ND | | Australia |
| MSMB2021WGS | ENV | 2a | GCF_001528425.1 | ND | | Australia |
| MSMB575WGS | ENV | 2a | GCF_001534105.1 | ND | | Australia |
| MSMB576WGS | ENV | 2a | GCF_001531955.1 | ND | | Australia |
| MSMB612WGS | ENV | 2a | GCF_001532145.1 | ND | | Australia |
| NCTC13007 | CF | 1 | GCF_900446205.1 | | LMG 13010 | Belgium |
| NKI379 | ENV | 2a | GCF_001302465.1 | ND | | Taiwan |
| R-20526 | ENV | 1 | GCF_001267755.1 | 836 | | Belgium |

### Table S2 Quality statistics of the 73 *B. multivorans* draft bacterial genomes sequenced in this study

| Strain | Size | No. of contigs | GC content (%) | Shortest contig size | Median sequence size | Mean sequence size | Longest contig size | N50 Value |
|---|---|---|---|---|---|---|---|---|
| BCC0006 | 6272301 | 107 | 67.3 | 119 | 1564 | 58619.6 | 923050 | 239280 |
| BCC0009 | 6342332 | 83 | 67.2 | 130 | 2945 | 76413.6 | 1042576 | 226583 |
| BCC0032 | 6506850 | 70 | 67.1 | 129 | 1739 | 92955 | 597776 | 396722 |
| BCC0033 | 6697341 | 82 | 67.1 | 101 | 1488 | 81674.9 | 873117 | 361065 |
| BCC0043 | 6544285 | 95 | 67.1 | 121 | 1132 | 68887.2 | 946841 | 383582 |
| BCC0047 | 7115961 | 80 | 66.6 | 122 | 2624 | 88949.5 | 1060012 | 360758 |
| BCC0065 | 6507164 | 64 | 67.4 | 101 | 1505 | 101674.4 | 731639 | 407234 |
| BCC0066 | 6510316 | 101 | 67.2 | 100 | 1564 | 64458.6 | 471548 | 225974 |
| BCC0068 | 6440630 | 197 | 67.1 | 101 | 1519 | 32693.6 | 377625 | 154108 |
| BCC0074 | 6541253 | 52 | 67.3 | 117 | 28069 | 125793.3 | 741944 | 401179 |
| BCC0075 | 6612635 | 78 | 67.2 | 103 | 11573 | 84777.4 | 938953 | 268072 |
| BCC0079 | 6991234 | 82 | 66.8 | 108 | 2155 | 85259 | 947587 | 337014 |
| BCC0080 | 6706461 | 113 | 66.8 | 101 | 1294 | 59349.2 | 686101 | 289154 |
| BCC0082 | 6621312 | 305 | 67.2 | 101 | 954 | 21709.2 | 304664 | 92055 |
| BCC0084 | 6599540 | 137 | 67.1 | 101 | 1857 | 48171.8 | 377138 | 165313 |
| BCC0087 | 6452471 | 62 | 67.4 | 105 | 4359 | 104072.1 | 859151 | 493463 |
| BCC0093 | 6278118 | 46 | 67.23 | 312 | 40751 | 143203.1 | 739761 | 275222 |
| BCC0096 | 6375796 | 91 | 67.2 | 109 | 2853 | 70063.7 | 1058392 | 233122 |
| BCC0101 | 6384309 | 113 | 67.1 | 120 | 2162 | 56498.3 | 665106 | 219505 |
| BCC0134 | 6414252 | 97 | 67.4 | 129 | 1574 | 66217.5 | 460055 | 308606 |
| BCC0141 | 6171198 | 83 | 67.4 | 110 | 16803 | 74351.8 | 689872 | 216985 |
| BCC0181 | 6537738 | 62 | 67.3 | 101 | 893 | 105554.1 | 835514 | 408687 |
| BCC0188 | 6652537 | 144 | 67 | 114 | 9369 | 46198.2 | 572214 | 164596 |
| BCC0225 | 6773647 | 59 | 67.1 | 127 | 1882 | 114807.6 | 1416856 | 6405560 |
| BCC0241 | 6655513 | 94 | 67.1 | 125 | 7260 | 70803.3 | 631291 | 200202 |
| BCC0246 | 6471839 | 65 | 67.3 | 122 | 16255 | 99566.8 | 678041 | 271451 |
| BCC0247 | 6531380 | 92 | 67.3 | 111 | 2137 | 70993.3 | 731592 | 286611 |
| BCC0255 | 6734567 | 87 | 67.2 | 108 | 1413 | 77497.6 | 510982 | 327645 |
| BCC0264 | 6629658 | 60 | 67.1 | 114 | 27268 | 110494.3 | 519753 | 365645 |
| BCC0266 | 6527276 | 92 | 67.3 | 115 | 12924 | 70948.7 | 447105 | 196973 |
| BCC0292 | 6360854 | 110 | 67.1 | 103 | 1675 | 57904.6 | 771986 | 207640 |
| BCC0293 | 6508922 | 198 | 67 | 102 | 1540 | 32956.4 | 537897 | 193063 |
| BCC0300 | 6582551 | 74 | 67.2 | 108 | 1428 | 89061.5 | 729256 | 386587 |
| BCC0303 | 6252321 | 96 | 67.2 | 18 | 8940 | 63799.2 | 707653 | 192319 |
| BCC0317 | 6355099 | 81 | 67.5 | 106 | 9257 | 78458 | 758055 | 235405 |
| BCC0321 | 6244833 | 106 | 67.32 | 102 | 2970 | 58975.8 | 659695 | 210819 |
| BCC0375 | 6289743 | 100 | 67.3 | 103 | 3795 | 62897.4 | 893830 | 253362 |
| BCC0381 | 6248359 | 82 | 67.3 | 134 | 3188 | 76199.5 | 725497 | 229512 |
| BCC0384 | 6378847 | 159 | 67.4 | 100 | 449 | 40118.5 | 485421 | 244802 |
| BCC0470 | 6984344 | 91 | 66.8 | 108 | 1413 | 76838.7 | 947619 | 337014 |
| BCC0493 | 6530376 | 126 | 67.1 | 109 | 722 | 51828.4 | 450290 | 261403 |
| BCC0497 | 6789721 | 97 | 66.9 | 121 | 2101 | 69997.1 | 592166 | 259787 |
| BCC0533 | 6410978 | 78 | 67.35 | 124 | 1058 | 82311.4 | 1043364 | 421975 |
| BCC0553 | 6694432 | 66 | 67.09 | 101 | 1625 | 101527.8 | 930655 | 405205 |
| BCC0583 | 7040902 | 286 | 66.73 | 101 | 2597 | 24693.6 | 426567 | 131205 |
| BCC0585 | 6447635 | 61 | 67.4 | 105 | 4145 | 105765.1 | 694113 | 337699 |
| BCC0702 | 6455034 | 110 | 67.1 | 108 | 3289 | 58682.1 | 515667 | 241164 |
| BCC0704 | 6449118 | 105 | 67.15 | 108 | 6604 | 61471.5 | 463159 | 217445 |
| BCC0710 | 6234591 | 89 | 67.5 | 100 | 887 | 70051.6 | 684558 | 310618 |
| BCC0729 | 6316285 | 139 | 67.31 | 105 | 1312 | 45512.6 | 468355 | 188298 |
| BCC0737 | 6223066 | 76 | 67.3 | 102 | 18360 | 81882.4 | 599007 | 239689 |
| BCC0814 | 6273978 | 110 | 67.3 | 101 | 2755 | 57036.2 | 427585 | 238427 |
| BCC0865 | 6791523 | 134 | 66.6 | 106 | 2148 | 50683 | 557856 | 262358 |
| BCC0901 | 6497146 | 61 | 67.17 | 13 | 1752 | 106579.9 | 866533 | 529217 |
| BCC0904 | 6029426 | 163 | 67.4 | 100 | 5437 | 36990.3 | 380474 | 106352 |
| BCC0907 | 6658191 | 99 | 66.93 | 124 | 1949 | 67349.9 | 839825 | 265179 |
| BCC0915 | 6277422 | 87 | 67.32 | 121 | 2755 | 72227.8 | 594460 | 243519 |
| BCC0921 | 6786237 | 158 | 66.6 | 106 | 2274 | 42950.9 | 557856 | 197853 |
| BCC0962 | 6277315 | 88 | 67.32 | 121 | 2972 | 71405.8 | 594459 | 243519 |

| | | | | | | | | |
|---|---|---|---|---|---|---|---|---|
| BCC0968 | 6785185 | 143 | 66.58 | 107 | 2303 | 47500.3 | 557857 | 231462 |
| BCC1147 | 6576406 | 109 | 67.1 | 105 | 5295 | 60334 | 565382 | 173107 |
| BCC1148 | 6573791 | 130 | 67.1 | 104 | 1652 | 50567.6 | 365455 | 171919 |
| BCC1177 | 6520943 | 160 | 67.1 | 101 | 2151 | 40755.9 | 470646 | 148928 |
| BCC1185 | 6383426 | 65 | 67.4 | 105 | 4145 | 98206.6 | 692368 | 368878 |
| BCC1190 | 6366933 | 112 | 67.1 | 109 | 6177 | 56847.6 | 378394 | 176650 |
| BCC1271 | 6997886 | 219 | 66.8 | 101 | 2320 | 32022.5 | 434582 | 151267 |
| BCC1272 | 6882314 | 163 | 66.8 | 117 | 3934 | 42222.8 | 772166 | 178859 |
| BCC1367 | 6246910 | 99 | 67.2 | 108 | 7439 | 63156.2 | 707677 | 207005 |
| BCC1368 | 6273724 | 109 | 67.2 | 102 | 2542 | 57557.1 | 796635 | 218602 |
| BCC1384 | 6246946 | 97 | 67.2 | 108 | 7439 | 64458.8 | 707668 | 220614 |
| BCC1385 | 6606699 | 112 | 66.9 | 101 | 1752 | 58988.4 | 544646 | 236600 |
| BCC1421 | 6609469 | 70 | 67.2 | 102 | 14809 | 94461.5 | 940166 | 352409 |
| **Average** | **6514719.4** | **107.09722** | **67.143056** | **109.81944** | **5156.1111** | **69942.958** | **676024.39** | **348751.403** |

**Table S3. PCR primer sequences designed for all four *B. multivorans* lineage-specific target genes**

| Target Gene | Primer Name | Primer Sequence (5' to 3') [c] | Primer Length (bp) | Position | Annealing Temperature (°C) | Product Size (bp) |
|---|---|---|---|---|---|---|
| ***YiaJ_1*** | YIAJBM1F | **AT**C**C**GG**C**AA**CT**ATTCGCT | 18 | 4007519-4007536[a] | 53.3 | 537 |
| | YIAJBM1R | **CAACGCTTTCCGTAGATG** | 18 | 4007000-4007017[a] | | |
| ***ghrB_1*** | GHRBBM1F | CAAGCAACCGACCGAA**AG** | 18 | 4008677-4008694[a] | 53.0 | 744 |
| | GHRBBM1R | GGAGACAG**A**ATCACGTT**C** | 18 | 4009403-4009420[a] | | |
| ***naiP_3*** | NAIPBM1F | **A**GCCGC**C**GA**AC**AA**A**G**A**TT**GA** | 20 | 4014235-4014254[a] | 55.7 | 981 |
| | NAIPBM1R | CTGAAGCCGGTCAGAAA**G** | 18 | 4015198-4015215[a] | | |
| ***glnM_2*** | GLNMBM2F | T**G**AA**T**G**CCG**GCCACGTA**TG** | 19 | 1792198-1792216[b] | 55.5 | 322 |
| | GLNMBM2R | GACGCATACGACAG**T**TCC | 18 | 1791895-1791912[b] | | |

[a] Position relative to the BCC0084 (lineage 1) complete genome

[b] Position relative to the ATCC 17616 (lineage 2a) complete genome

[c]Mismatches for each primer sequence are highlighted in bold. Red indicates a mismatch in all strains of the opposing lineage and blue indicates mismatches within target lineage strains

**Table S4. Growth kinetics of the *B. multivorans* strains showing growth rate, lag phase and maximum OD**

| Isolate | Statistical Model | Growth Rate ($h^{-1}$) | Lag Phase (λ) | Maximum OD |
|---|---|---|---|---|
| ATCC 17616 | richards | 0.066393 | 5.142631 | 0.32093 |
| BCC0006 | richards | 0.02309 | 5.307211 | 0.359725 |
| BCC0009 | gompertz | 0.044397 | 2.467438 | 0.375609 |
| BCC0032 | richards | 0.019392 | 1.586864 | 0.349948 |
| BCC0033 | logistic | 0.035193 | 4.217101 | 0.369231 |
| BCC0043 | richards | 0.024118 | 4.811215 | 0.373581 |
| BCC0047 | richards | 0.02602 | 4.995805 | 0.394584 |
| BCC0065 | logistic | 0.037451 | 4.248356 | 0.369135 |
| BCC0246 | gompertz | 0.043649 | 3.473084 | 0.411566 |
| BCC0303 | richards | 0.020057 | 11.59027 | 0.351278 |
| BCC0497 | richards | 0.016688 | 7.899568 | 0.35352 |
| BCC0269 | richards | 0.021785 | 10.1249 | 0.327047 |
| BCC0737 | richards | 0.031135 | 7.56969 | 0.373028 |
| BCC0302 | gompertz | 0.045548 | 3.481668 | 0.383461 |
| BCC0225 | gompertz | 0.018589 | 2.508097 | 0.326467 |
| BCC1177 | richards | 0.020631 | 4.565362 | 0.337437 |
| BCC0084 | richards | 0.033811 | 3.66836 | 0.382379 |
| BCC1185 | richards | 0.020373 | 9.216607 | 0.319333 |
| BCC0074 | richards | 0.053576 | 4.486977 | 0.358003 |
| BCC0710 | richards | 0.035296 | 4.439582 | 0.349044 |
| BCC0266 | richards | 0.037115 | 3.610816 | 0.39047 |
| BCC0080 | richards | 0.037799 | 4.237065 | 0.397279 |
| BCC0493 | richards | 0.02316 | 12.34905 | 0.368741 |
| BCC0066 | logistic | 0.045924 | 4.124916 | 0.37575 |
| BCC0865 | richards | 0.018893 | 7.541105 | 0.341761 |
| BCC0068 | richards | 0.014588 | 7.214126 | 0.331623 |
| BCC0096 | richards | 0.041078 | 3.280156 | 0.37302 |
| BCC0247 | richards | 0.014531 | 0.7697 | 0.373392 |
| BCC0921 | richards | 0.020943 | 12.20387 | 0.333387 |
| BCC0082 | richards | 0.03056 | 3.437512 | 0.372343 |
| BCC0317 | gompertz | 0.042476 | 2.43051 | 0.380466 |
| BCC0141 | richards | 0.020178 | 4.394878 | 0.305782 |
| BCC1148 | richards | 0.034645 | 5.522386 | 0.401968 |
| BCC0814 | gompertz | 0.016017 | 2.446835 | 0.394281 |
| BCC0264 | logistic | 0.040676 | 3.827443 | 0.373854 |
| BCC0904 | richards | 0.04133 | 3.591944 | 0.394376 |
| BCC0075 | richards | 0.016353 | 4.902274 | 0.312914 |
| BCC0375 | richards | 0.019472 | 3.500513 | 0.36421 |
| BCC0087 | richards | 0.042158 | 3.400263 | 0.38021 |
| BCC0702 | richards | 0.016862 | 1.362989 | 0.335749 |
| BCC0101 | richards | 0.048303 | 3.296633 | 0.380578 |
| BCC0079 | richards | 0.029847 | 3.006941 | 0.366436 |
| BCC0241 | logistic | 0.04607 | 3.650837 | 0.372825 |
| BCC0384 | gompertz.exp | 0.043328 | 3.879821 | 0.355711 |
| BCC0188 | richards | 0.018039 | 1.466231 | 0.357657 |
| BCC1147 | gompertz.exp | 0.04965 | 4.450516 | 0.335572 |
| BCC1190 | richards | 0.040893 | 2.944074 | 0.371125 |
| BCC1272 | richards | 0.059492 | 4.869471 | 0.327952 |
| BCC1368 | richards | 0.028289 | 3.644625 | 0.364084 |
| BCC1385 | richards | 0.044277 | 2.76252 | 0.362452 |

### Table S5. Swimming and swarming motility within the *B. multivorans* strains

| Strain | Mean motility zone (mm) within each growth medium and motility phenotype [b,c] | | |
|---|---|---|---|
| | 0.3% LB (Swimming) | 0.5% LB (Swarming) | 0.5% BSM-G (Swarming) |
| ATCC 17616 [a] | 21.8 | 7.8 | 22.2 |
| BCC0006 | 10.3 | 0.0 | 0.0 |
| BCC0009 | 57.8 | 14.4 | 90.0 |
| BCC0032 | 18.8 | 2.6 | 3.8 |
| BCC0033 | 25.2 | 12.0 | 9.8 |
| BCC0043 | 15.3 | 2.8 | 0.0 |
| BCC0047 | 25.3 | 10.0 | 30.1 |
| BCC0065 | 24.1 | 11.8 | 8.8 |
| BCC0066 | 22.3 | 10.4 | ND |
| BCC0068 | 0.0 | 0.0 | 0.0 |
| BCC0074 | 20.1 | 3.8 | 3.1 |
| BCC0075 | 41.2 | 17.3 | 77.3 |
| BCC0079 | 29.4 | 18.8 | ND |
| BCC0080 | 26.8 | 3.5 | 0.0 |
| BCC0082 | 31.8 | 9.9 | ND |
| BCC0084 | 58.6 | 24.3 | 36.3 |
| BCC0087 | 27.3 | 17.4 | 24.6 |
| BCC0096 | 38.8 | 12.3 | 18.1 |
| BCC0101 | 23.7 | 5.3 | 8.8 |
| BCC0141 | 13.9 | 8.3 | 13.6 |
| BCC0188 | ND | ND | ND |
| BCC0225 | 30.1 | 11.7 | ND |
| BCC0241 | 29.5 | 16.4 | 18.0 |
| BCC0246 | 18.7 | 5.4 | 5.8 |
| BCC0247 | 13.8 | 5.3 | 6.5 |
| BCC0264 | 21.3 | 4.7 | 3.5 |
| BCC0266 | 25.1 | 10 | 11.1 |
| BCC0269 | 8.5 | 5.4 | 4.0 |
| BCC0302 | 46.1 | 14.3 | ND |
| BCC0303 | 2.3 | 2.7 | 14.8 |
| BCC0317 | 90.0 | 19.5 | 87.5 |
| BCC0375 | 19.8 | 11.8 | 9.0 |
| BCC0381 | ND | ND | ND |
| BCC0384 | 17.3 | 5.1 | 10.8 |
| BCC0493 | 6.8 | 4.5 | 0.0 |
| BCC0497 | 5.3 | 2.8 | 16.2 |
| BCC0702 | 47.1 | 32.6 | 69.1 |
| BCC0710 | 22.3 | 8 | 6.3 |
| BCC0737 | 9.9 | 3.8 | 11.8 |
| BCC0814 | 24.4 | 14.3 | 16.8 |
| BCC0865 | 7.8 | 5.7 | 4.3 |
| BCC0904 | 3.2 | 2 | 0.0 |
| BCC0921 | 10.7 | 7.5 | 3.3 |
| BCC1147 | 24.8 | 8.3 | 26.6 |
| BCC1148 | 14.7 | 5.3 | 14.3 |
| BCC1177 | 60.9 | 24.6 | 90.0 |
| BCC1185 | 17.3 | 9.5 | 1.2 |
| BCC1190 | 59.6 | 17.8 | 90.0 |
| BCC1272 | 25.8 | 7.3 | 70.7 |
| BCC1368 | 7.7 | ND | ND |

| | | | |
|---|---|---|---|
| BCC1385 | 25.6 | 14.3 | 40.4 |

[a] *B. multivorans* ATCC 17616 was used as a positive motile control for all assays
[b] Highly motile strains (≤50 mm average motility diameter) are highlighted in blue
[c] ND = mean not determined due to variable results

### Table S6. Colony morphology and EPS production of *B. multivorans*

| Lineage | Strain | Morphology | | | | |
|---|---|---|---|---|---|---|
| | | Shape | Size | Appearance | Congo Red Agar | EPS production (YEM agar) |
| 1 | BCC0006 | Round | Small | Shiny | Red | - |
| | BCC0009 | Irregular | Medium | Shiny | Pink to pink with orange centre | ++++ |
| | BCC0010 | Round | Small | Shiny | Pink with red centre | ++++ |
| | BCC0067 | Round | Small | Shiny | Pink with red centre | ++++ |
| | BCC0080 | Round | Medium | Shiny | Pink with red centre | ++ |
| | BCC0084 | Irregular | Medium | Shiny | Pink with orange centre | +++ |
| | BCC0093 | Round | Small | Shiny | Pink with red centre | ++++ |
| | BCC0101 | Round | Medium | Shiny | Pink with red centre | ++ |
| | BCC0115 | Irregular | Medium | Shiny | Pink with orange centre | ++++ |
| | BCC0141 | Round | Small | Shiny | pink with red centre | ++++ |
| | BCC0149 | Round | Small | Shiny | Pink with orange-red centre | ++++ |
| | BCC0175 | Round | Medium | Shiny | Pink with orange-red centre | ++ |
| | BCC0292 | Round | Medium | Shiny | Pink with red centre | ++++ |
| | BCC0293 | Round | Small | Shiny | Pink with orange centre | ++ |
| | BCC0303 | Round | Small | Shiny | Pink | +++ |
| | BCC0321 | Round | Small | Shiny | Pink | + |
| | BCC0375 | Round | Medium | Shiny | Pink with red centre | ++ |
| | BCC0702 | Round | Small | Shiny | Pink with orange centre | +++ |
| | BCC0704 | Round | Small | Shiny | Pink with orange centre | +++ |
| | BCC0737 | Round | Small | Shiny | Pink with red centre | +++ |
| | BCC0814 | Round | Medium | Shiny | Pink to pink with orange centre | +++ |
| | BCC0865 | Round | Small | Shiny | Pink with red centre | ++ |
| | BCC0904 | Round | Small | Shiny | Pink | + |
| | BCC0915 | Round | Small | Shiny | Pink with orange-red centre | +++ |
| | BCC0921 | Round | Small | Shiny | Pink with orange centre | + |
| | BCC0962 | Round | Small | Shiny | Pink with red centre | ++++ |
| | BCC0968 | Round | Small | Shiny | Pink with orange-red centre | +++ |
| | BCC1177 | Round | Small | Shiny | Pink with orange centre | +++ |
| | BCC1190 | Round | Small | Shiny | Pink with red centre | ++++ |
| | BCC1367 | Round | Small | Shiny | Orange to red | ++ |
| | BCC1384 | Round | Small | Shiny | Pink | ++++ |
| | BCC1385 | Irregular | Medium | Shiny | Pink with red centre | ++++ |
| 2 | ATCC17616 | Irregular | Medium | Shiny | Pink with red centre | +++ |
| | BCC0005 | Round | Medium | Shiny | Pink with red centre | ++++ |
| | BCC0008 | Round | Small | Shiny | Pink with orange-red centre | ++++ |
| | BCC0031 | Round | Medium | Shiny | Pink with orange-red centre | ++++ |
| | BCC0032 | Round | Medium | Shiny | Pink with red centre | ++ |
| | BCC0033 | Irregular | Medium | Shiny | Pink with red centre | +++ |
| | BCC0037 | Round | Medium | Shiny | Pink with orange-red centre | ++++ |
| | BCC0043 | Round | Small | Shiny | Pink with orange centre | ++ |
| | BCC0047 | Round | Medium | Shiny | pink | ++++ |
| | BCC0050 | Round | Medium | Shiny | Pink with orange-red centre | ++++ |
| | BCC0059 | Round | Small | Shiny | Orange to red | +++ |
| | BCC0065 | Round | Medium | Shiny | Pink to pink with orange centre | ++ |
| | BCC0066 | Round | Medium | Shiny | Pink with red centre | ++ |
| | BCC0068 | Round | Small | Shiny | pink | - |
| | BCC0074 | Round | Medium | Shiny | Pink with red centre | ++ |
| | BCC0075 | Round | Small | Shiny | Pink with orange centre | ++++ |
| | BCC0079 | Round | Medium | Shiny | Pink | ++++ |
| | BCC0082 | Irregular | Medium | Shiny | Pink with red centre | ++ |
| | BCC0087 | Round | Medium | Shiny | pink with orange centre | ++ |
| | BCC0089 | Round | Medium | Shiny | Pink with orange-red centre | ++++ |
| | BCC0096 | Round | Medium | Shiny | Pink with orange centre | +++ |
| | BCC0099 | Round | Medium | Shiny | Pink with red centre | +++ |
| | BCC0102 | Round | Small | Shiny | Pink with red centre | ++++ |
| | BCC0134 | Round | Small | Shiny | Pink with red centre | ++ |
| | BCC0181 | Round | Small | Shiny | Pink with red centre | ++++ |
| | BCC0188 | Round | Small | Shiny | Pink with red centre | - |
| | BCC0225 | Irregular | Medium | Shiny | Red | +++ |
| | BCC0241 | Round | Medium | Shiny | pink with red centre | ++ |
| | BCC0244 | Round | Medium | Shiny | Pink with red centre | + |
| | BCC0246 | Round | Medium | Shiny | Pink with red centre | ++++ |
| | BCC0247 | Round | Medium | Shiny | Pink with red centre | ++ |
| | BCC0255 | Round | Small | Shiny | Pink with orange centre | ++++ |
| | BCC0264 | Round | Medium | Shiny | Pink with red centre | + |
| | BCC0266 | Round | Medium | Shiny | pink with red centre | ++ |
| | BCC0269 | Round | Medium | Shiny | pink with red centre | +++ |
| | BCC0300 | Round | Small | Shiny | Pink with orange centre | +++ |
| | BCC0317 | Irregular | Medium | Shiny | Pink with orange centre | ++++ |
| | BCC0384 | Round | Small | Shiny | pink | + |
| | BCC0470 | Round | Medium | Shiny | Pink with red centre | +++ |
| | BCC0493 | Round | Small | Shiny | Pink | - |
| | BCC0497 | Round | Small | Shiny | Pink | - |

| | | | | | | |
|---|---|---|---|---|---|---|
| | BCC0533 | Irregular | Medium | Shiny | Pink with red centre | ++++ |
| | BCC0553 | Round | Small | Shiny | Pink with orange-red centre | + |
| | BCC0583 | Round | Small | Shiny | Pink with red centre | ++ |
| | BCC0585 | Round | Small | Shiny | Pink with red centre | +++ |
| | BCC0585 | Round | Small | Shiny | Pink with orange-red centre | +++ |
| | BCC0710 | Round | Small | Shiny | Pink with red centre | ++ |
| | BCC0729 | Irregular | Medium | Shiny | Pink with red centre | +++ |
| | BCC0901 | Round | Small | Shiny | Red | +++ |
| | BCC0907 | Round | Medium | Shiny | Pink with red centre | +++ |
| | BCC1147 | Round | Small | Shiny | Pink with orange centre | ++ |
| | BCC1148 | Round | Small | Shiny | pink | ++ |
| | BCC1185 | Round | Small | Shiny | Pink | +++ |
| | BCC1271 | Round | Small | Shiny | Pink with orange centre | +++ |
| | BCC1272 | Round | Small | Shiny | Pink with red centre | +++ |
| | BCC1421 | Round | Small | Shiny | Pink with orange-red centre | ++++ |
| Other | BCC1368 | Round | Small | Shiny | Red | + |

### Supplementary Figures

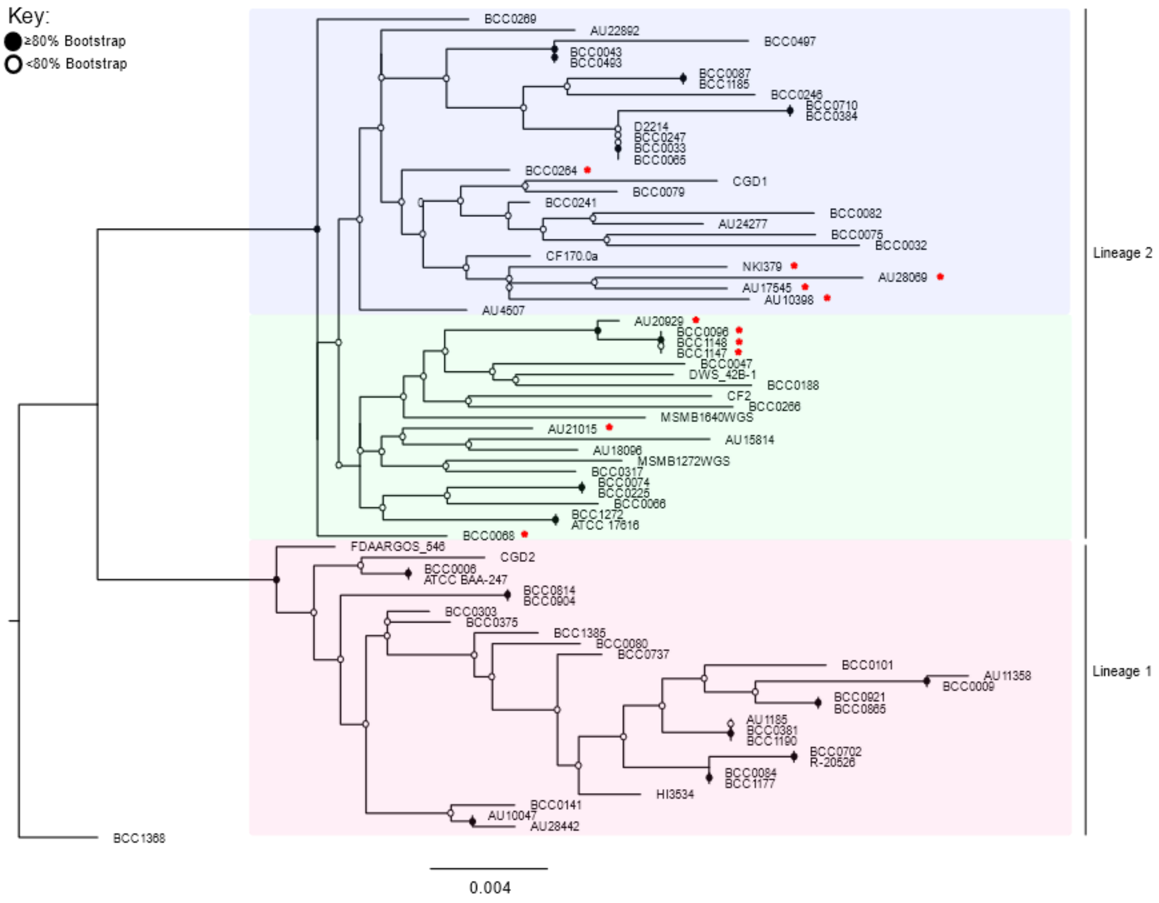


**Figure S1. MLST-gene phylogeny of the 77 genome B. multivorans strain panel.** The RAxML (100 bootstraps) phylogenetic tree was created by extracting the MLST sequences from the sequenced whole genomes using MLST check (Page et al. 2016b). Bootstrap values are indicated using the key. Filled circles represent a bootstrap of ≥80% and a hollow (white) circle means confidence of ≤80%. Lineages are shown on the right hand side of the figure, with colour coding illustrating the lineages and sub-lineages. Lineage 1 = red, lineage 2a = green, lineage 2b = blue. The red asterix next to strain names represents the flipped genomes in lineage 2. Scale bar represents the phylogenetic distance of 0.004 nucleotide substitutions.

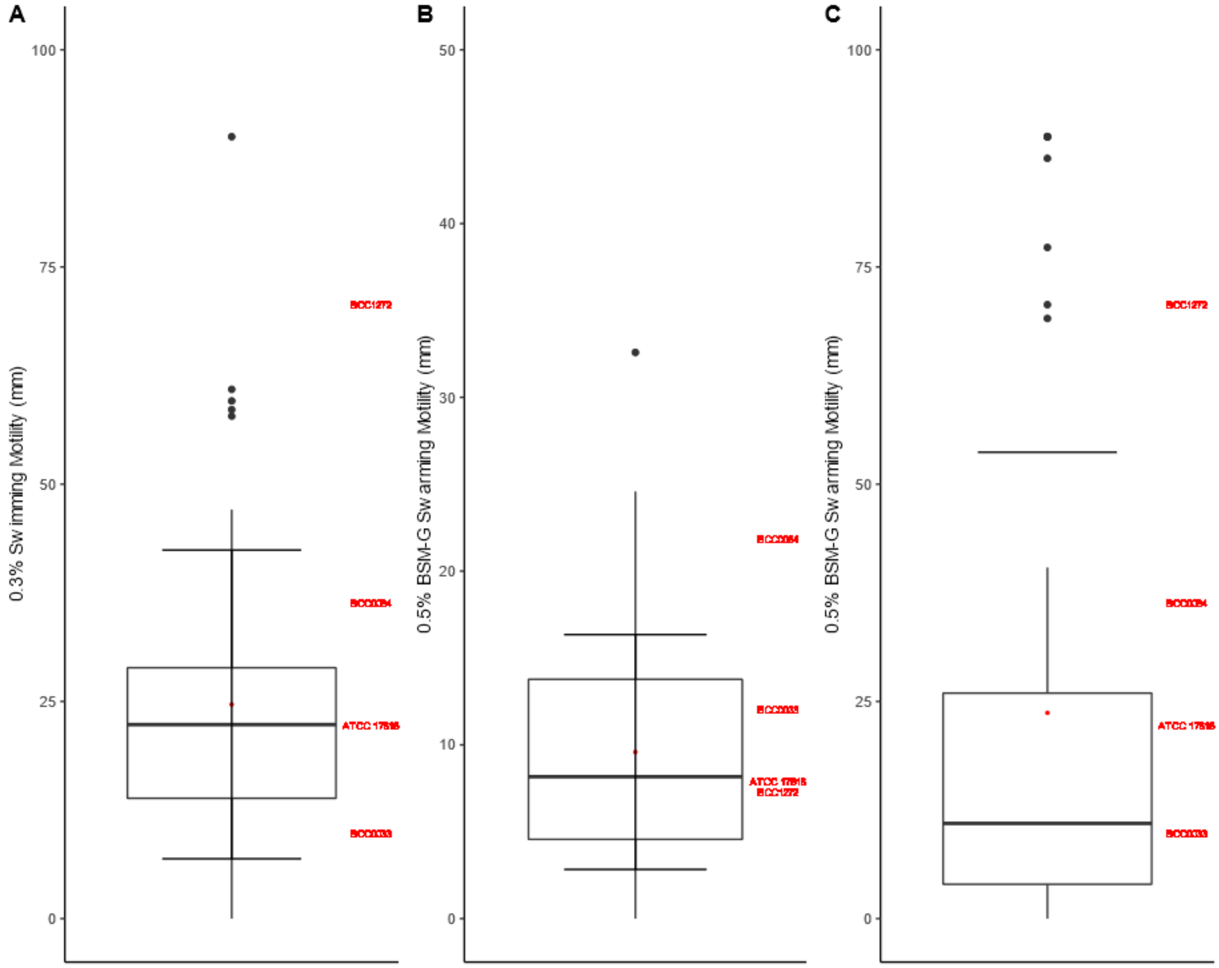


**Figure S2. Swimming and swarming motility in the B. multivorans strain panel after 24-hours incubation at 37°C.** (A) Swimming motility was performed on 0.3% LB agar (n = 49). Swarming motility was performed on (B) 0.5% LB agar (n = 48) and (C) 0.5% BSM-G agar (n = 44). Model *B. multivorans* strains (n = 4) are annotated in red on the righthand side of each box plot. No motility associations statistically linked to each *B. multivorans* lineage were identified.

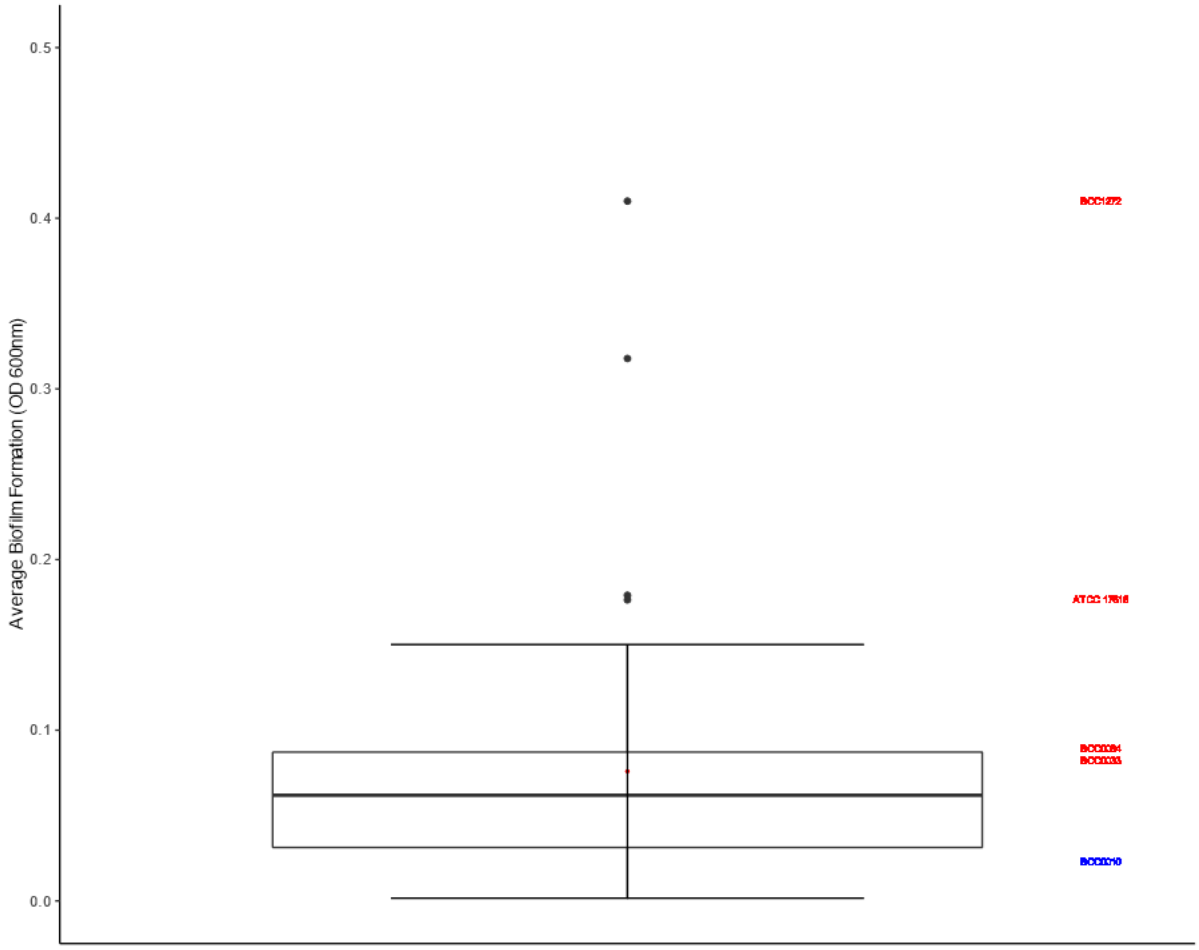


**Figure S3. Biofilm formation of the *B. multivorans* strain panel (n = 49) after 24-hours.** Among of biofilm was assessed using the crystal violet assay, reading the results using a plate reader at $600_{nm}$. Biofilm controls were *B. multivorans* ATCC 17616 for the 'high' former and BCC0010 (blue) for the 'low' control. Model CF B. multivorans strains are noted in red on the right-hand side of the box plot. Box plots show the mean, lower quartile, and upper quartile for biofilm staining. Outliers are also shown. No biofilm formation abilities statistically linked to each *B. multivorans* lineage were identified.
